# Repeat turnover meets stable chromosomes: repetitive DNA sequences mark speciation and gene pool boundaries in sugar beet and wild beets

**DOI:** 10.1101/2023.09.01.555723

**Authors:** Nicola Schmidt, Katharina Sielemann, Sarah Breitenbach, Jörg Fuchs, Boas Pucker, Bernd Weisshaar, Daniela Holtgräwe, Tony Heitkam

## Abstract

**Background:** Sugar beet (*Beta vulgaris* subsp. *vulgaris*) and its crop wild relatives share a base chromosome number of nine and similar chromosome morphologies. Yet, interspecific breeding is impeded by chromosome and sequence divergence that is still not fully understood. Since repetitive DNA sequences represent the fastest evolving parts of the genome, they likely impact genomic variability and contribute to the separation of beet gene pools. Hence, we investigated if innovations and losses in the repeatome can be linked to chromosomal differentiation and speciation.

**Results:** We traced genome- and chromosome-wide evolution across sugar beet and twelve wild beets comprising all sections of the beet genera *Beta* and *Patellifolia*. For this, we combined data from short and long read sequencing, flow cytometry, and cytogenetics to build a comprehensive data framework for our beet panel that spans the complete scale from DNA sequence to chromosome up to the genome.

Genome sizes and repeat profiles reflect the separation of the beet species into three gene pools. These gene pools harbor repeats with contrasting evolutionary patterns: We identified section- and species-specific repeat emergences and losses, e.g. of the retrotransposons causal for genome expansions in the section *Corollinae/Nanae*. Since most genomic variability was found in the satellite DNAs, we focused on tracing the 19 beetSat families across the three beet sections/genera. These taxa harbor evidence for contrasting strategies in repeat evolution, leading to contrasting satellite DNA profiles and fundamentally different centromere architectures, ranging from chromosomal uniformity in *Beta* and *Patellifolia* species to the formation of patchwork chromosomes in *Corollinae/Nanae* species.

**Conclusions:** We show that repetitive DNA sequences are causal for genome size expansion and contraction across the beet genera, providing insights into the genomic underpinnings of beet speciation. Satellite DNAs in particular vary considerably among beet taxa, leading to the evolution of distinct chromosomal setups. These differences likely contribute to the barriers in beet breeding between the three gene pools. Thus, with their isokaryotypic chromosome sets, beet genomes present an ideal system for studying the link between repeats, genome variability, and chromosomal differentiation/evolution and provide a theoretical basis for understanding barriers in crop breeding.

## Background

### Interplay between chromosomal stability and genome evolution

Large evolutionary leaps in terms of speciation and genome divergence are occasionally accompanied by chromosome number changes caused by polyploidisation events and/or structural rearrangements (*Arabidopsis*: Yogeeswaran *et al*., 2005; *Carex*: Escudero *et al*., 2012 & 2015; *Cylicomorpha*: Rockinger *et al*., 2016). In contrast, some genera display genomic divergence despite harboring karyotypes with stable base chromosome numbers across all known species (McCann *et al*., 2020; Vitales *et al*., 2020a; Pellicer *et al*., 2021). That even extends to large intraspecific differences in genome size despite a consistent chromosome number (*Euphrasia*: Becher *et al*., 2021). Therefore, the question arises how inter- and intraspecific genomic divergence accumulates while the chromosomal setup is maintained.

One of the links between the genome and its partitioning into chromosomes are repetitive DNA sequences, which provide structure to eukaryotic karyotypes. Despite their fast evolution, repetitive DNA sequences have a conserved function by providing sequence material to the main structural regions, such as centromeres and telomeres. Nevertheless, how the evolution of the repeatome itself relates to the global chromosome and genome evolution is still a matter of debate (e.g. centromere paradox; Henikoff *et al*., 2001; Presting, 2018).

### Species of the Beta and Patellifolia genera are marked by dynamic genomes, but stable chromosome numbers

We wondered how genomic diversity despite stability of chromosomal number is reflected in the repeat composition. To test this, we used species of the beet genera *Beta* and *Patellifolia* as examples – plant taxa that have been used to study repetitive DNA evolution for over thirty years (Schmidt *et al*., 1990; and later publications from the lab of late Thomas Schmidt). The members of these two genera belong to chromosomally stable taxa, all marked by a base chromosome number of x=9 (despite some higher ploidies) and containing roughly equally-sized, metacentric chromosomes.

The genera *Beta* and *Patellifolia* comprise at least eleven species that are separated by up to 38.4 million years of evolution (Hohmann *et al*., 2006). Species from the genus *Beta* are divided into two or three different sections (*Beta*, *Corollinae/Nanae*) depending on whether the endemic species *B. nana* is considered a separate section (Kadereit *et al*., 2006; Frese and Ford-Lloyd, 2020; Sielemann *et al*., 2022). Cultivated beets such as sugar beet are varieties of *B. vulgaris* subsp. *vulgaris* within the section *Beta*. The presumed progenitor of all cultivated beets, *B. vulgaris* subsp. *maritima*, also belongs to this section (Frese and Ford-Lloyd, 2020). For better readability, *B. vulgaris* subsp. *vulgaris*, *B. vulgaris* subsp. *adanensis*, and *B. vulgaris* subsp. *maritima* will be hereafter referred to as *B. vulgaris*, *B. adanensis*, and *B. maritima*, respectively.

Sugar beet is a relatively young crop that went through an exceptionally narrow bottleneck during its 200 years of domestication (Fischer, 1989). This has led to very low genetic diversity and a loss of several valuable traits such as pathogen resistances and tolerance to adverse environmental conditions (e.g. drought, salinated soil; Panella *et al*., 2020). Therefore, it is necessary to harvest genetic variation in the beet germplasm to improve cultivated beet varieties. However, crossing experiments of sugar beet with many of its crop wild relatives (CWRs) resulted in reproduction-defective offspring such as sterile and semi-fertile, as well as aneuploid and anorthoploid plants indicating that postzygotic isolation mechanisms, e.g. the lack of chromosome homology due to larger genomic differences, are causal for the limited gain of genetically improved seeds rather than prezygotic isolation mechanisms (Frese and Ford-Lloyd, 2020).

Based on the crossability with *B. vulgaris*, the wild beets are grouped into three different gene pools. These correspond to the three beet main taxa with section *Beta* species representing the primary gene pool, *Corollinae/Nanae* species representing the secondary gene pool, and *Patellifolia* species representing the tertiary gene pool. Given the fact that di- and polyploid species are present in all of these gene pools (Frese and Ford-Lloyd, 2020) and due to the chromosome uniformity across all beet clades, polyploidisation and restructuring of chromosomes seem to play a rather subordinate role in the emergence of genomic variety in *Beta* and *Patellifolia* species. Instead, differences at the DNA sequence level (Sielemann *et al*., 2023a) may be the cause of flawed chromosome pairing, thus resulting in the observed crossing barriers.

The rapid evolution of repetitive elements can be a major factor in reduced pairing between homologous and homoeologous chromosomes (Dvorak, 1983) as it leads to varying repeat compositions in the genomes of even closely related species. Repetitive DNA sequences include transposable elements (TEs) and tandem repeats (TRs) that differ in their origin, amplification mode, and sequence characteristics (Bennetzen, 2005; Bennetzen and Wang, 2014).

There are two classes of TEs: class I or retrotransposons with a ‘copy-and-paste’ transposition mechanism and an RNA intermediate, and class II or DNA transposons with a ‘cut-and-paste’ mechanism, thus DNA being the intermediate in replication. Class I is divided into two subclasses, those flanked by long terminal repeats (LTR retrotransposons) and those without (non-LTR retrotransposons; Finnegan, 1989; Wells and Feschotte, 2020). The classes and subclasses are further divided hierarchically into order, superfamily, family, subfamily, and lineage (as reviewed in Wicker *et al*., 2007; Piégu *et al*., 2015; Neumann *et al*., 2019).

TRs on the other hand include ribosomal genes, telomeres and satellite DNAs. Satellite DNAs (satDNAs) are highly repetitive non-coding sequences that are arranged in large tandem arrays and contribute to structurally important chromosomal regions such as the centromeres (Lower *et al*., 2018; Garrido-Ramos, 2021).

Beet genomes, in particular the sugar beet genome, are well characterized regarding retrotransposons (Wollrab *et al*., 2012; Weber *et al*., 2013; Heitkam *et al*., 2014; Schwichtenberg *et al*., 2016; Maiwald *et al*., 2021), DNA transposons (Menzel *et al*., 2006; Menzel *et al*., 2012), and even related viral relics (Schmidt *et al*., 2021). Additionally, at least 13 satDNA families are characterized (e.g. Dechyeva and Schmidt, 2006; Zakrzewski *et al*., 2010). In beets, the most iconic repeat identified so far is the centromeric satDNA pBV (Schmidt and Metzlaff, 1991), which differs from the centromeric satDNA in more distant wild beets (pTS5, pTS4.1; Schmidt and Heslop-Harrison, 1996; Gindullis *et al*., 2001). However, there is not yet a full picture of the repeat landscape across beet genomes that is needed to identify the evolutionary relationships across time and across species. Expanding the knowledge about the repeat landscapes may help to explain the different crossabilities between the beet gene pools. To understand the impact of repeats on crossing barriers, beet genomic resources are needed. So far there are three published reference genome sequence assemblies for sugar beet at our disposal (consecutive RefBeet versions: Dohm *et al*., 2014; consecutive EL10 versions: Funk *et al*., 2018 and McGrath *et al*., 2020; KWS2320ONT_v1.0: Sielemann *et al*., 2023b) and the number of studies comparing the wild beet germplasm to cultivated varieties is continuously increasing (Sielemann *et al*., 2022, 2023a; Wascher *et al*., 2022). Hence, sugar beet and its CWRs are well-suited to serve as a reference for maintaining stable karyotypes despite global changes across the repetitive genome.

To illuminate how the karyotypically stable wild beet genomes differ from another, we focused on repetitive DNA sequences as one of the most impactful contributors to genomic variability. We asked how repeat evolution across wild beets can be linked to speciation and, hence, to crossing barriers. We further investigated if the wild beet germplasm’s splits into species, sections, and genera (i.e. primary, secondary, and tertiary gene pools) are mirrored by broad genomic innovations in the repeatome. For this, we chose 17 beet accessions, including all major beet taxa, measured their genome sizes, and determined their chromosome numbers and karyotypes. To add a (pan-) genomic layer, we generated low-pass whole genome shotgun data for all accessions as well as long reads for selected beet genomes. After assessing the phylogenetic placements of these accessions (Sielemann *et al*., 2022, 2023a), we estimated their repetitive DNA content and finely classified their repetitive DNA sequences in the respective TE and TR hierarchies. This not only allows following repetitive DNA evolution comprehensively across the beet genera, determining repeat gains, losses and replacements, but also linking it back to chromosomal location and karyotypic stability.

## RESULTS

### Corollinae species have the largest monoploid genome sizes among sugar beet and wild beets

To provide the foundation for later analyses of genome and chromosome evolution, we first determined the chromosome configuration and ploidy levels of all 17 cultivated and wild beet accessions, and then estimated their genome sizes by flow cytometry (Additional file 1: Fig. S1; Table 1). Building onto the plastome-based phylogenetics framework (Sielemann *et al*., 2022), our 17-beet-species-panel allows comprehensive investigation of the *Beta* and *Patellifolia* germplasm.

**Table 1:** Sampled beet and wild beet species, including five-letter code, accession details, ploidy level / chromosome number, and genome size. Gene pools are marked according to Frese and Ford-Lloyd, 2020.

| beet gene pools | species | code | accession | ENA accession no. | chromosome configuration | genome size 1C [Mbp] |
| --- | --- | --- | --- | --- | --- | --- |
| primary | *Beta vulgaris* subsp. *vulgaris* | Bvul1 | KWS2320 | ERR6110425 | 2n=2x=18 | 712 |
|  | *Beta vulgaris* subsp. *adanensis* | Bada1 | BETA 1233 | ERR6110427 | 2n=2x=18 | 709 |
|  | *Beta vulgaris* subsp. *maritima* | Bmar1 | BETA 1101 | ERR6110429 | 2n=2x=18 | 703 |
|  | *Beta vulgaris* subsp. *maritima* | Bmar2 | BETA 2322 | ERR6110431 | 2n=2x=18 | 694 |
|  | *Beta patula* | Bptu1 | BETA 548 | ERR6110433 | 2n=2x=18 | 706 |
|  | *Beta macrocarpa* | Bmca1 | BETA 881 | ERR6110435 | 2n=4x=36 | 1370 |
| secondary | *Beta lomatogona* | Blom1 | BETA 674 | ERR6110449 | 2n=2x=18 | 929 |
|  | *Beta macrorhiza* | Bmrh1 | BETA 830 | ERR6110451 | 2n=2x=18 | 936 |
|  | *Beta macrorhiza* | Bmrh2 | BETA 576 | ERR6110453 | 2n=2x=18 | 946 |
|  | *Beta corolliflora* | Bcor1 | BETA 408 | ERR6110441 | 2n=4x=36 | 2027 |
|  | *Beta intermedia* | Bint1 | BETA 431 | ERR6110455 | 2n=5x=45 | 2468 |
|  | *Beta intermedia* | Bint2 | BETA 923 | ERR6110457 | 2n=5x=45 | 2471 |
|  | *Beta nana* | Bnan1 | BETA 546 | ERR6110459 | 2n=2x=18 | 766 |
|  | *Beta nana* | Bnan2 | BETA 570 | ERR6110461 | 2n=2x=18 | 754 |
| tertiary | *Patellifolia procumbens* | Ppro1 | BETA 951 | ERR6110463 | 2n=2x=18 | 712 |
|  | *Patellifolia webbiana* | Pweb1 | BETA 526 | ERR6110465 | 2n=2x=18 | 723 |
|  | *Patellifolia patellaris* | Ppat1 | BETA 892 | ERR6110467 | 2n=4x=36 | 1502 |

The beet genome sizes are in accordance with the respective ploidy level (Table 1; Additional file 1: Fig. S1). Diploid members of the section *Beta* have 1C genomes sizes of approx. 700 Mbp, whereas the genome of the sole tetraploid species within this section, *B. macrocarpa*, is nearly twice as large. In contrast, diploid members of the section *Corollinae* achieve considerably higher values with an average 1C genome size of almost 940 Mbp. Also, the polyploid species *B. corolliflora* and *B. intermedia* have genome sizes slightly larger than mathematically expected (2027 Mbp instead of 1880 Mbp, and approx. 2470 Mbp instead of 2350 Mbp). With a value of approx. 760 Mbp, the genome size of the diploid *B. nana* is more similar to that of the diploids from the section *Beta* than from the section *Corollinae*. Regarding the sister genus, diploid *Patellifolia* species have genome sizes similar to that of the diploids from the section *Beta*, whereas the genome of the tetraploid *P. patellaris* is again slightly larger than mathematically expected (1502 Mbp instead of 1435 Mbp).

### Repeat abundance and genome size correlate positively for all major repeat types, except for tandem repeats

We estimated repeat proportions in the genomes of all species through individual as well as comparative clustering using the RepeatExplorer2 pipeline. Combined, the repeats identified for each species represent between 53% (Bptu) and 68% (Bint) of the total genome (Additional file 1: Table S1). For better comparability and to minimize the dependence of the genome size from the ploidy level, polyploid species were also analyzed with downsampled read sets so that they correspond to a diploid chromosome set (‘ploidy corrected genome size’). In general, we observe a high correlation between repeat proportion (absolute values) and genome size, with r2 = 0.996 (p < 8.94e-17) for the ploidy corrected genome sizes (Fig. 1A). Further, we observe a clustering of species with larger genome sizes and higher repeat contents (*Corollinae* members) vs. species with smaller genome sizes and lower repeat contents (section *Beta*, *B. nana*, and the genus *Patellifolia*; Fig. 1A).

**Fig. 1:**
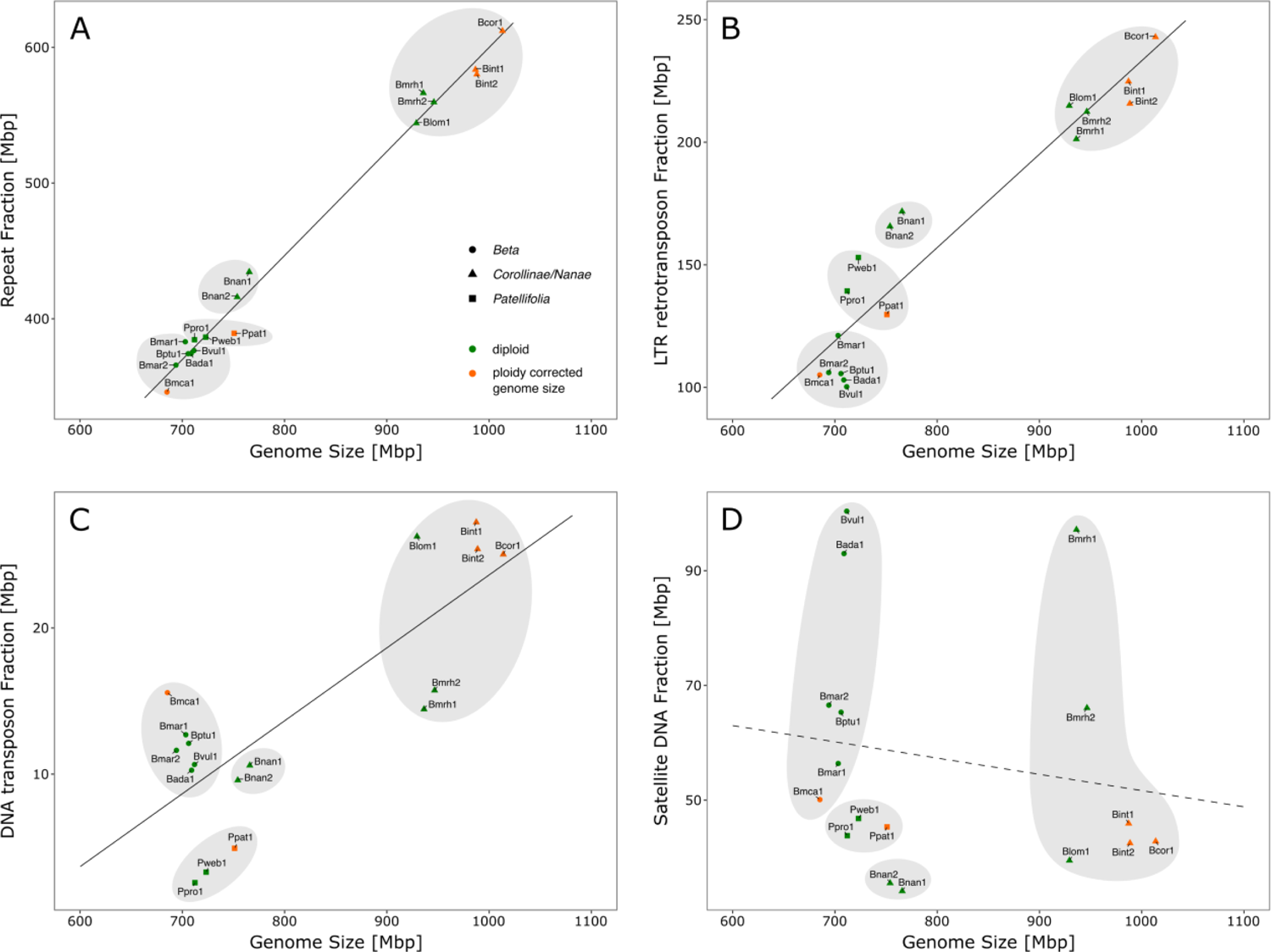
Correlation between repeat fraction and genome size in *Beta* and *Patellifolia* species. For better comparability with natural diploids (green), polyploid species were downsampled as artificial diploids (orange). Shapes and shades indicate the different beet sections and genera, respectively. Samples are abbreviated according to Table 1. (A) Positive correlation between the total repeat fraction and genome size. (B) Positive correlation between LTR retrotransposon fraction and genome size. (C) Positive correlation between DNA transposon fraction and genome size. (D) The satellite DNA fraction and genome size tend to relate negatively.

When individual repeat classes (i.e. LTR retrotransposons, DNA transposons, and satDNAs) are considered, the distinct taxonomic groups can be resolved according to their genome sizes and respective repeat contents (absolute values; Fig. 1B-D, shadings). Again, a positive correlation can be observed when the overall proportion of LTR retrotransposons (r2 = 0.948, p = 7.32e-9) and DNA transposons (r2 = 0.798, p = 1.22e-4) is plotted against the ploidy corrected genome sizes, respectively (Fig. 1B, C). However, when specifically focusing on the proportion of satDNAs, an extraordinarily high amount (up to roughly 14%, see Additional file 1: Table S1) is found in the beet species with small genome sizes, in particular *B. vulgaris* and *B. adanensis*, whereas the beet species with the largest genome sizes from the section *Corollinae/Nanae* (with the exception of *B. macrorhiza*) show the lowest contents of satDNAs (less than 5%). This results in a negative trend regarding the relation between the satDNA proportion and the genome size (r2 = −0.166, p > 0.52; Fig. 1D). The same observations apply for the overall and individual repeat fractions calculated as proportions (Additional file 1: Fig. S2).

### Genomic TEs differences across sugar beet and wild beets reflect the separation into beet sections and genera

The read clustering results based on a 0.1× coverage of the ploidy corrected genome size for each species are analyzed. The repetitive fraction of all analyzed genomes is composed mainly of LTR retrotransposons (14-30% of the genome) with twice the relative content in *B. corolliflora* and *B. intermedia* (*Corollinae* section) compared to species of the section *Beta* (see Additional file 1: Table S1). With the exception of *B. vulgaris*, in which Ty1-*copia* and Ty3-*gypsy* retrotransposons show a rather balanced relation to each other, in most other genomes, the Ty3-*gypsy* elements dominate (Fig. 2A-C, see Additional file 1: Table S1). This observation is most pronounced in *P. webbiana*, where the proportion of Ty3-*gypsy* retrotransposons is four times bigger than that of Ty1-*copia* retrotransposons.

**Fig. 2:**
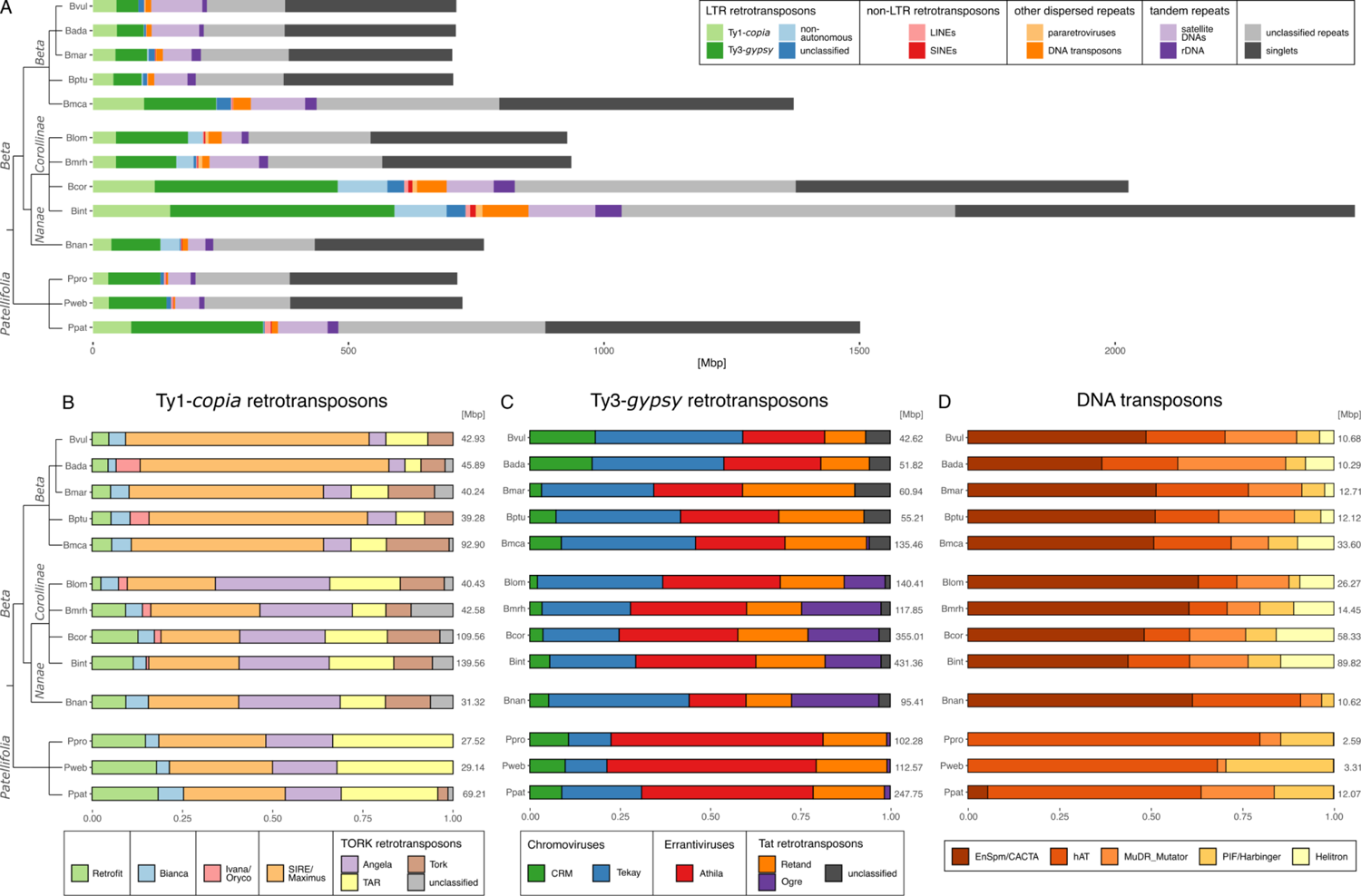
Repeat composition of several *Beta* and *Patellifolia* species. The analyzed samples and abbreviations accord to accession 1 of the respective species (see Table 1) as the repeat compositions of different accessions from the same species are highly similar (see Additional file 1: Table S1). (A) Proportion of major repeat classes shown in relation to the total genome size. (B) Composition of the Ty1-*copia* LTR retrotransposon fraction. (C) Composition of the Ty3-*gypsy* LTR retrotransposon fraction. (D) Composition of the DNA transposon fraction.

As a substantial difference between species from the section *Beta* and the remaining taxa (section *Corollinae/Nanae* and genus *Patellifolia*), their most abundant LTR retrotransposons belong to different superfamilies: In section *Beta*, SIRE/Maximus retrotransposons from the Ty1-*copia* superfamily contribute the highest TE share (3-4%), whereas the genomes of the other species consist largely of Athila retrotransposons from the Ty3-*gypsy* superfamily (4-6% in *Corollinae* species; 8-9% in *Patellifolia* species). However, *B. nana* stands out as the proportion of Athila elements (2%) within its genome is clearly surpassed by the proportion of Ogre (3%) and Tekay elements (5%). In this respect, *B. nana* resembles the species from the section *Beta* in which Tekay elements are the most abundant Ty3-*gypsy* retrotransposons as well (2-4% of the genome, with approximately as many Tekay elements as SIRE/Maximus elements in the tetraploid *B. macrocarpa*).

Another notable observation is that beets from the section *Corollinae/Nanae* are characterized by a quite high amount of non-autonomous LTR retrotransposons (3-5%, see Additional file 1: Table S1) compared to the other species. The investigation of the corresponding clusters revealed that this increased share can be traced back to one single, so far uncharacterized element.

Taking a look at the DNA transposon fraction, species from the genus *Beta* are dominated by EnSpm/CACTA terminal inverted repeat (TIR) transposons, whereas the most abundant DNA transposons in *Patellifolia* species belong to the hAT superfamily (Fig. 2D). We find a higher genome proportion of DNA transposons in species with larger genome sizes, with *Patellifolia* species generally having fewer DNA transposons compared to beets of the genus *Beta*.

### TEs are generally conserved across the beet genomes and show section-specific abundances

An all-to-all comparison across the beets (accession 1 of every species; see Table 1) revealed that the overall most abundant repeat, a so far unknown LTR retrotransposon of the Athila/Errantivirus lineage, occurs in all analyzed genomes, representing a genome proportion of 0.89% (Bnan1) up to 8.08% (Pweb1; Additional file 1: Fig. S3). However, the high Athila/Errantivirus proportion in *Corollinae* species is caused by another LTR retrotransposon originally described as five distinct dispersed repeats by Gao *et al*. (2000; pBC1054, pBC227, pBC305, pBC507/169, and pBC537). In general, about half of the repeat families are found in all analyzed beet genomes. The other half consists of species-, section-, and genus-specific repeats. There are fewer *Patellifolia*-specific repeats (33 out of 395 clusters) than *Beta*-specific repeats (172 out of 395 clusters), resulting in a lower overall repeat diversity in the genus *Patellifolia*. However, the *Patellifolia*-specific repeats are highly abundant, whereas all repeats in the *Beta* genomes (with the exception of some satDNAs, see paragraph below) are rather moderately abundant. Yet, we also find section-specific repeats with regard to the sections *Beta* (8 out of 172 genus Beta-specific clusters) and *Corollinae/Nanae* (11 out of 172 genus *Beta*-specific clusters). Several LTR retrotransposons (Athila, Tekay, Tat, and Ale/Retrofit elements) seem to be re-amplified in the beet genomes of the *Corollinae/Nanae* section, whereas an increased abundance of SIRE/Maximus elements was observed in the beet genomes of the *Beta* section. Furthermore, a so far unknown Ogre LTR retrotransposon, as well as the non-autonomous LTR retrotransposon mentioned in the paragraph above, set *Corollinae/Nanae* apart from all other analyzed beets (see Fig. 2C and Additional file 1: Fig. S3).

### Beet satDNAs break ranks (1): Beta and Patellifolia satDNAs are few, but highly amplified, whereas Corollinae/Nanae satDNAs are diverse and lowly abundant

Comparing *Beta* and *Patellifolia* genomes, the greatest repeatome difference is found in the satDNAs (Fig. 3). They contribute roughly 6% to the genomes of *Patellifolia* species and only 4% to the genomes of the *Corollinae/Nanae* species (with the exception of *B. macrorhiza*; see Additional file 1: Table S1). In contrast, in genomes of the section *Beta*, satDNAs contribute up to 14%, which is about as much as the LTR retrotransposon fraction.

**Fig. 3:**
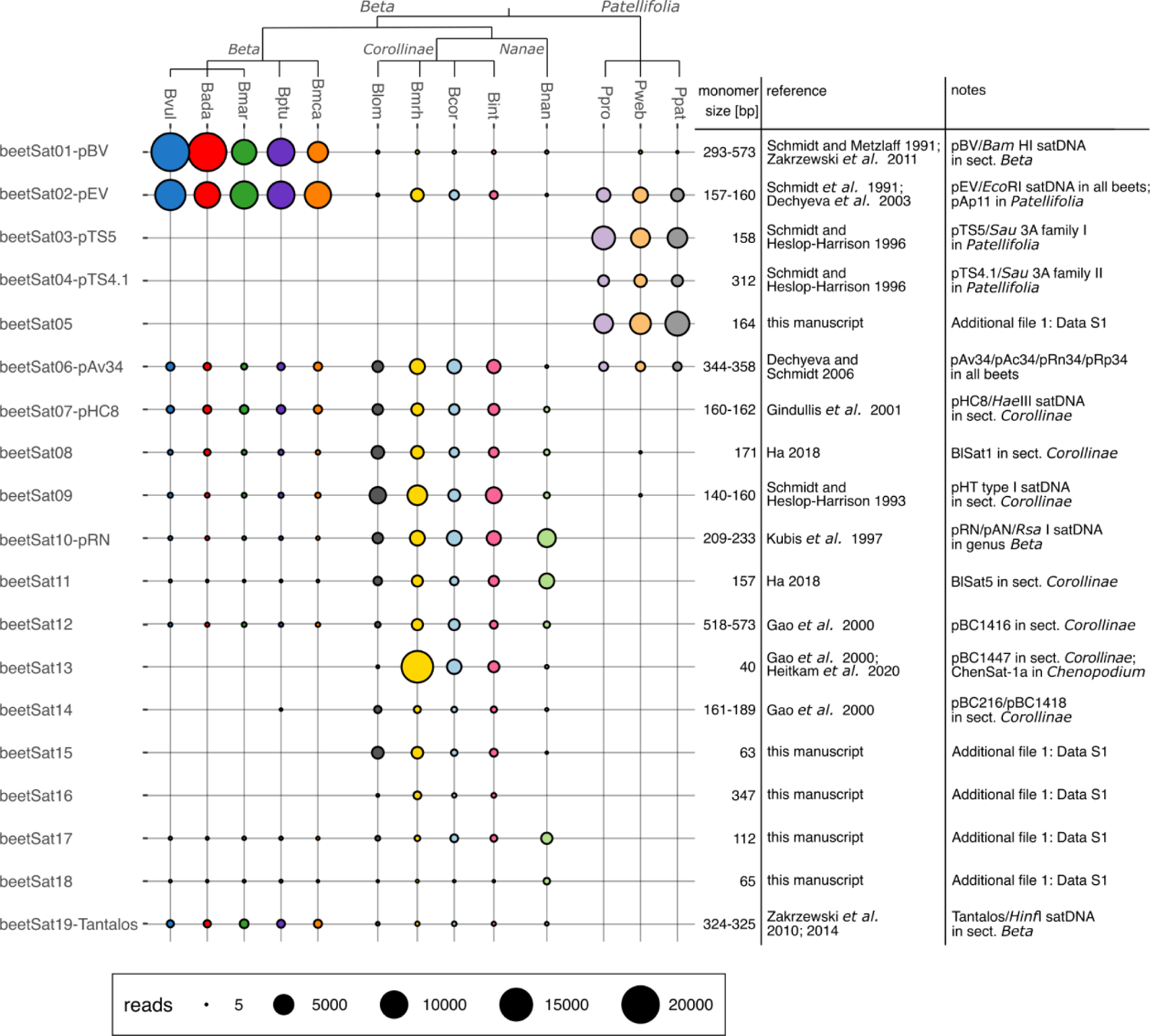
Quantification of satDNAs within different *Beta* and *Patellifolia* genomes. The relative satDNA abundance is indicated by the size of the circles, which corresponds to the number of short reads within the respective satDNA cluster. The monomer sizes as well as relevant references are provided. Further previously described ‘satDNAs’ and minisatellites (i.e. Dione/*Fok*I-satellite and Niobe/*Alu*I-satellite: Zakrzewski et al. 2010, 2014; BvMSats: Zakrzewski et al. 2010; BvuSats: Li et al. 2021) were not included as we found that these repeats are not arranged in long arrays and therefore we consider them tandem repeats rather than satDNA. The monomer sequences of newly identified satDNAs can be found in Additional file 1: Data S1.

The reason for this divergence is the extraordinarily high abundance of two iconic sugar beet satDNAs (see Additional file 1: Fig. S3), the centromeric beetSat01-pBV and the intercalary beetSat02-pEV (Schmidt and Metzlaff, 1991; Schmidt *et al*., 1991). BeetSat01-pBV and beetSat02-pEV are highly abundant in all species of the section *Beta* contributing up to 8% and 5% of the genomes, respectively (Fig. 3; see Additional file 1: Table S2). However, beetSat02-pEV also occurs in beets of the section *Corollinae/Nanae* and within the sister genus *Patellifolia* with low or moderate abundance (Fig. 3). In contrast, canonical beetSat01-pBV satDNA arrays (as described by Schmidt and Metzlaff, 1991; Zakrzewski *et al*., 2011) are strictly limited to the beets of the section *Beta*.

Although genomes of the *Patellifolia* genus contain less satDNAs than those of the section *Beta*, they are also dominated by abundant satDNA families (Fig. 3; see Additional file 1: Table S2): Again, the most prominent satDNAs in the *Patellifolia* species are those that constitute the centromeres: the centromeric beetSat03-pTS5 and the pericentromeric beetSat04-pTS4.1 (Schmidt & Heslop-Harrison, 1996). However, in sequence and monomer length they are substantially different from beetSat01-pBV. In addition, we identified another highly abundant and genus-specific satDNA (beetSat05). The satDNA designated as beetSat06 is the only satDNA that is distributed quite equally among all beet genomes (Fig. 3; Additional file 1: Table S2). This subtelomeric satDNA ubiquitously occurs on all chromosomes and was previously described under different names depending on the respective plant species (e.g. in *B. vulgaris* it is known as pAv34; Dechyeva & Schmidt, 2006). However, Dechyeva and Schmidt (2006) found that, in the wild beet *B. nana*, this satDNA is restricted to one single pair of chromosomes which is consistent with the reduced abundance of beetSat06 that we observed in *B. nana* using the comparative repeat analysis (Fig. 3).

Generally, within the *Corollinae/Nanae*, the specific satDNA quantities set *B. nana* slightly apart from the other species within this section (Fig. 3; Additional file 1: Table S2). This applies for other known satDNAs such as beetSat07-pHC8 (Gindullis *et al*., 2001; Fig. 4G, H) or beetSat10-pRN (Kubis *et al*., 1997; Fig. 4I, J), as well as for the satDNAs newly identified during the comparative repeat analysis (beetSat15–18; Additional file 1: Data S1 and Data S2). These satDNAs are most pronounced in beet genomes of the section *Corollinae/Nanae* without the conspicuously high abundances observed in the section *Beta* and the sister genus *Patellifolia*. Among the satDNAs that define the *Corollinae/Nanae* genomes, only beetSat13 is prominent, especially in *B. macrorhiza* (Fig. 3). This satDNA is also known as pBC1447 in *B. corolliflora* and ChenSat-1a in the related *Chenopodium quinoa* (Gao *et al*., 2000; Heitkam *et al*., 2020). Overall, regarding the interplay of satDNA diversity and abundance, we conclude that *Beta* and *Patellifolia* contain relatively few satDNA families that can reach high copy numbers. In contrast, species within the *Corollinae/Nanae* accumulate a wide satDNA variety with only low amplification levels (Fig. 3).

**Fig. 4:**
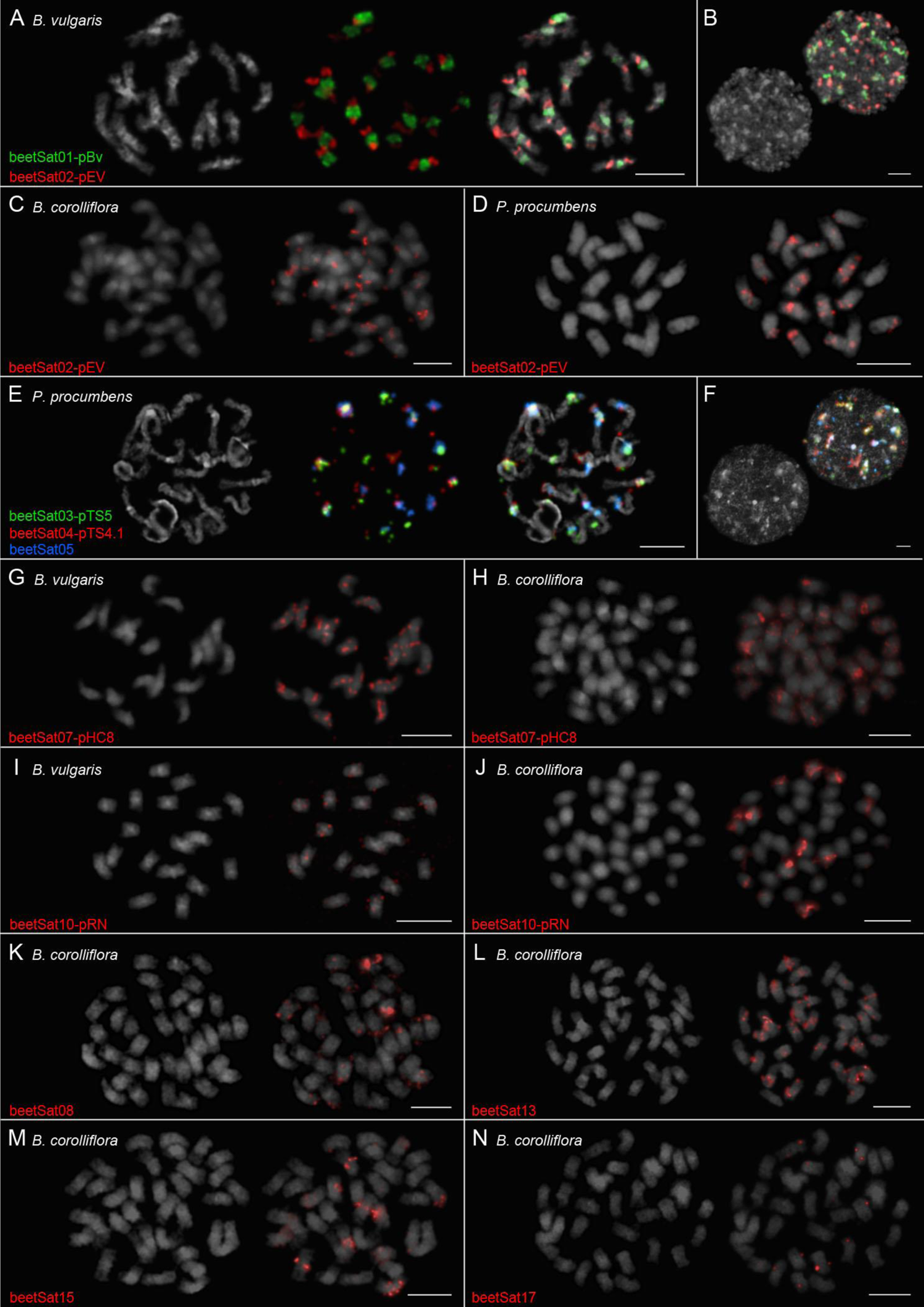
Multicolor fluorescent *in situ* hybridization to chromosome spreads of *B. vulgaris*, *B. corolliflora*, and *P. procumbens*. DAPI-stained mitotic chromosomes and interphase nuclei are shown in gray. (A) The centromeric and intercalary localization of the satDNAs beetSat01-pBV (green signals) and beetSat02-pEV (red signals), respectively, become visible on prometaphase chromosomes of *B. vulgaris*. (B) The interphase nucleus of *B. vulgaris* shows that beetSat01-pBV (green signals) and beetSat02-pEV (red signals) are located in heterochromatic regions. (C, D) The satDNA beetSat02-pEV can also be found on almost all chromosomes of *B. corolliflora* (C) and *P. procumbens* (D) in intercalary as well as distal regions, forming large as well as small arrays. Whereas the signals are mostly restricted on one chromosome arm each in *B. corolliflora* (C), they can be often found on both chromosome arms in *P. procumbens* (D). (E) The centromeric and pericentromeric localization of the satDNAs beetSat03-pTS5 (green signals) and beetSat04-pTS4.1 (red signals), respectively, become visible on prometaphase chromosomes of *P. procumbens*. Most centromeres (14 out of 18) also show large beetSat05 arrays (blue signals); in particular those that are not constituted by beetSat03-pTS5. (F) The interphase nucleus of *P. procumbens* shows that beetSat03-pTS5 (green signals), beetSat04-pTS4.1 (red signals), and beetSat05 (blue signals) are located in heterochromatic regions. (G, H) The satDNA beetSat07-pHC8 was detected on almost all arms of all chromosomes of *B. vulgaris* (G) and *B. corolliflora* (H): Signals were present in intercalary as well as distal regions, forming large as well as small arrays. Moreover, two centromeric beetSat07-pHC8 signals were detected in both species. (I, J) Hybridization sites for beetSat10-pRN are few in *B. vulgaris* (I), whereas large signal clusters were detected on most chromosomes of *B. corolliflora* (J). (I) The low beetSat10-pRN abundance in *B. vulgaris* is revealed by faint signals mostly on both arms of the respective chromosome. (J) The major hybridization sites of the beetSat10-pRN probe on *B. corolliflora* chromosomes are associated with the centromeres. Intercalary hybridization sites can be found mostly on one arm of further chromosomes. (K) The centromere of four *B. corolliflora* chromosomes showed signals (two major, two minor) after hybridization with the beetSat08 probe, whereas almost all other chromosomes showed intercalary and distal signals. (L) BeetSat13 can also be found at the centromeres of at least seventeen *B. corolliflora* chromosomes (ten major and seven minor signals). Eight further chromosomes hybridized in intercalary regions. (M) The beetSat15 hybridization resulted in at least six *B. corolliflora* chromosomes with signals at the centromeres and again eight chromosomes with intercalary signals. (N) BeetSat17 signals were detected on twelve *B. corolliflora* chromosomes with at least two of them being near the centromere. Information on probe labeling and detection can be found in the methods section. Scale bars = 5 µm.

### Beet satDNAs break ranks (2): Species- and section-specific array expansions lead to the emergence of unique satDNA landscapes across the beet genomes

As presented above, individual satDNAs can occur in high abundance in one beet species and in low abundance in another. To understand, if beet satDNA organization is retained across species and abundance patterns, we investigated their tandem arrangement across the three main beet taxa. By using long reads from *B. vulgaris* (section *Beta*), *B. corolliflora* (section *Corollinae*), and *P. procumbens* (genus *Patellifolia*) we showed that nearly all identified satDNAs occur in long tandem arrays in at least one of the three beet taxa (see dotplot visualization in Additional file 1: Fig. S4). Only beetSat18 does not show any long arrays in neither of the three species. This is easily explained as this satDNA is specific for *B. nana*, with head-to-tail arrangements identified on *B. nana* short reads.

We noted variation in monomer length and sequence (Additional file 1: Fig. S5), often with section specificity. Sugar beet’s main satDNA beetSat01-pBV is especially intriguing: It occurs only scarcely in *Patellifolia* genomes, forming no tandem arrays. However, in *B. corolliflora*, the beetSat01-pBV monomer is part of a different, tandemly arranged repeat, resulting in a much longer monomer size of approx. 1800 bp (Additional file 1: Fig. S4). This larger tandem repeat shows no further sequence similarity to publicly available nucleotide or protein database entries. Indeed, it may serve as a starting point to understand formation of one of the largest satDNA families in any plant genome. Similar potential start points of satDNA emergence were detected for beetSat10, beetSat11, beetSat12, beetSat17, and beetSat18 (Additional file 1: Fig. S4; arrows), suggesting a general pattern.

Further, monomer length variations were detected, indicating section specificity, but none as striking as for beetSat1-pBV. For instance, beetSat02-pEV variants diverge slightly in monomer length and sequence among the three beet taxa (Additional file 1: Fig. S5), whereas beetSat06 shows variation in higher order arrangement.

Overall, regarding satDNA amplification and array formation, we conclude that different satDNAs amplify in different beet species. Whereas large arrays can form in one wild beet genome, the same satDNA can occur only as a relic in the next. Similarly, emergence of species-/section-specific satDNA variants occurs. We observe that the respective beet satDNA landscapes are characterized by different evolutionary mechanisms: Whereas few *Beta* and *Patellifolia* satDNAs underwent amplification and homogenization, the satDNAs landscape of *Corollinae/Nanae* species is still rather dynamic, mirrored by the emergence of several new satDNAs.

### Beet satDNAs break ranks (3): Beta and Patellifolia satDNAs constitute major parts of all chromosomes, whereas Corollinae/Nanae satDNAs are restricted to chromosome subsets

To understand the structural chromosome makeup across the beets, we again focused on the three species *B. vulgaris*, *B. corolliflora* and *P. procumbens* to represent the three main beet taxa. First, we investigated the five main satDNAs: beetSat01-pBV to beetSat05.

Our two-color FISH onto sugar beet chromosomes shows that beetSat01-pBV (green) constitutes the centromeres, whereas beetSat02-pEV (red) builds large intercalary blocks along all *B. vulgaris* chromosomes (Fig. 4A, B; Kubis *et al*., 1998). In contrast, beetSat02-pEV is restricted to only a subset of chromosomes and/or comparatively small intercalary regions along *B. corolliflora* and *P. procumbens* chromosomes (Fig. 4C, D). This indicates that beetSat02-pEV is not only specific in its monomer sequence (see above), but also in its chromosomal localization among the three beet taxa.

In *P. procumbens*, beetSat03-pTS5 marks most, but not all centromeres (Fig. 4E, green signals; see also Gindullis *et al*., 2001). BeetSat05 resides at as many chromosomes (10-14 centromeres; Fig. 4E, blue signals). Thus, each centromere contains either beetSat03-pTS5 or beetSat05, or both. All centromeres are flanked by beetSat04-pTS4.1 that accompanies the centromeric beetSat03-pTS5 and beetSat05 signals, but also occupies some distal locations (Fig. 4E, red signals). All five main satDNAs are restricted to the DAPI-positive heterochromatin in interphase nuclei (Fig. 4B, F). Concluding, we note that chromosomes of *B. vulgaris* and *P. procumbens* – despite their phylogenetic distance – are organized similarly, with main satDNAs building the centromeres and intercalary regions.

This is in sharp contrast to the chromosome organization in *B. corolliflora*, a representative of the *Corollinae/Nanae* section: Out of the many satDNAs with varying degrees of amplification and homogenization (see above), we selected six for hybridizations (beetSat07-pHC8, beetsSat08, beetSat10-pRN, beetSat13, beetSat15, and beetSat17; Figure 4G-N). None of them is exclusively localized at the centromeres. Instead, intercalary signals are common as well and (peri-)centromeric signals are restricted to only some chromosomes, often being less prominent than centromeric *Beta/Patellifolia* signals. The comparison of two of these satDNAs at *B. vulgaris* and *B. corolliflora* chromosomes reveals that the signal patterns are more (beetSat07-pHC8) or less (beetSat10-pRN) similar in both genomes (Fig. 4G-J):

BeetSat07-pHC8 resides on all *B. vulgaris* chromosomes in intercalary and distal regions, but also close to at least two centromeres (Fig. 4G). In *B. corolliflora*, beetSat07-pHC8 resides in two centromeric and several intercalary regions as well. Major signals are pronounced on six chromosomes (presumably three chromosome pairs; Fig. 4H), including the two centromeric signals.

The second satDNA, beetSaat10-pRN, is lowly abundant in *B. vulgaris* (see Fig. 3), thus producing only few, faint and scattered signals (Fig. 4I). In contrast, beetSat10-pRN hybridized strongly to 6-8 chromosomes of *B. corolliflora*, predominantly in the centromeres (Fig. 4J).

To better understand how the patchy satDNA emergence/amplification patterns in the *Corollinae/Nanae* affect the chromosomes, we also localized beetSat08, beetSat13, beetSat15, and beetSat17 (Figure 4K-N):

BeetSat08 resides on nearly all *B. corolliflora* chromosomes (Fig. 4K). Two major and two minor signals mark the centromeres of four chromosomes, whereas the majority of signals was rather weak and distributed over the intercalary and distal regions.

BeetSat13, which is known to be associated with the centromeres of *C. quinoa* chromosomes (ChenSat-1a; Heitkam *et al*., 2020), can also be found at the centromeres of at least ten *B. corolliflora* chromosomes (Fig. 4L). BeetSat13 signals near the centromere were detected on seven additional chromosomes. The remaining chromosomes did not hybridize with the beetSat13 probe or showed intercalary beetSat13 signals.

All six major sites of beetSat15 hybridization are associated with *B. corolliflora* centromeres as well (Fig. 4M).

The comparatively low abundance of beetSat17 (see Fig. 3) resulted in only few hybridization signals in *B. corolliflora* (Fig. 4N): Signals were detected on twelve chromosomes with at least two of them showing weak centromeric signals.

Overall, regarding the chromosomal impacts of the vastly different satDNA landscapes in *Beta* and *Patellifolia* versus *Corollinae/Nanae*, we note that the different evolutionary patterns affect chromosome structure, especially the centromeres. *Beta* and *Patellifolia* centromeres are made up of few, highly abundant, homogenized satDNAs. Instead, the *Corollinae/Nanae* harbor ‘patchwork centromeres’: We identified at least 28 (peri-)centromeric signals with six different satDNAs (beetSat07-pHC8, beetSat08, beetSat10-pRN, beetSat13, beetSat15, beetSat17) on *B. corolliflora* chromosomes. From this, we conclude that *Corollinae/Nanae* centromeres are constituted by a multitude of different satDNAs, whereas *Beta* and *Patellifolia* centromeres are constituted by one or two main satDNAs, respectively.

## DISCUSSION

### A beet species panel to understand the evolving genome under karyotypic stability

We leverage a comprehensive repeatome study across 17 accessions of cultivated and wild beet species, spanning two sister genera, *Beta* and *Patellifolia*. We target all major species across all sections, including higher polyploids. Building onto the plastome-based phylogenetics framework (Sielemann *et al*., 2022, 2023a), our 17-beet-species-panel represents a well-characterized sugar beet and CWR panel with regard to tracing genome evolution, chromosome stability, pangenomics, and taxonomy.

### Beet and wild beet genome size variation results from repeat content fluctuations

Considering the respective ploidy level, the beet genome sizes show relatively moderate variation: 1.4-fold variation among the diploids, 1.5-fold variation among the tetraploids, and also 1.5-fold variation among the ploidy corrected genome sizes. Meanwhile, intraspecific differences are minimal, with almost no variation. For taxa with high chromosomal variability, e.g. across *Euphrasia* individuals (1.3-fold variation), structural changes such as the loss or gain of chromosome fragments are likely responsible for genome size variations (Becher *et al*., 2021). In contrast, we hypothesize that genome size variation within plant taxa with stable chromosome setups, such as the analyzed beet species, depends strongly on genomic repeats. This can be demonstrated by a clear correlation between genome size and repeat fraction. For the beet species, we have found such a correlation not only between the overall repeat content, but especially between the LTR retrotransposons and the respective genome size, indicating that the amplification and elimination of LTR retrotransposons in particular is causal for genome size differences between beets of the same ploidy. In comparison to other angiosperms, the determined correlation is at least as high (e.g. compared to *Fabeae* sp.; Macas *et al*., 2015), if not higher (e.g. compared to *Eleocharis* sp., *Solanum* sp., *Hesperis*-clade sp.; Zedek *et al*., 2010; Michael, 2014; Gaiero *et al*., 2019; Hloušková *et al*., 2019; Gantuz *et al*., 2021), pointing to a particularly strong impact of repeats (i.e. LTR retrotransposons) on beet genome sizes.

Specific amplification and elimination of repeats may explain why the genome size of *B. nana* rather resembles those of the section *Beta*, even though this species is repeatedly considered a member of the section *Corollinae* (Kadereit *et al*., 2006; Frese and Ford-Lloyd, 2020; Sielemann *et al*., 2022). However, genome sizes of beets from the genus *Patellifolia* are most similar to those of the section *Beta*, suggesting that repeat amplification and/or acquisition caused a genome upsizing in *Corollinae/Nanae* species.

### Repeat abundances mirror the phylogenies of the beet and wild beet sections

With 53-68%, the repetitive fraction of beet genomes represents typical values compared to other Amaranthaceae (approx. 76% in quinoa: Heitkam *et al*., 2020; approx. 51% in spinach: Li *et al*., 2021). Discrepancies in repeat abundances to a previous RepeatExplorer2 analysis in *B. vulgaris* (Kowar *et al*., 2016) are moderate and result from the exclusion of organellar DNA from our read set as well as the improvement of the repeat annotation by including a more comprehensive, custom repeat database. In general, the repeat content results not only from the number of repetitive sequences, but also from their type: A repeat with a longer element structure accounts for a larger share of the genome than a shorter repeat. Within the beet genomes, the most frequent repeats belong to the LTR retrotransposons, namely the Ty3-*gypsy* retrotransposons, which include the longest known TEs in plants (i.e. Ogre elements with >23 kb in length; Orozco-Arias *et al*., 2019). The predominance of Ty3-*gypsy* retrotransposons within the repeat fraction was observed for other angiosperms as well (Kelly *et al*., 2015; Macas *et al*., 2015; Gaiero *et al*., 2019; Hloušková *et al*., 2019; Dodsworth *et al*., 2020). However, the rather balanced share of Ty3-*gypsy* and Ty1-*copia* retrotransposons in *B. vulgaris* may reflect different TE dynamics in the domesticated beet cultivars, whereas the independent amplification and/or acquisition of Ty3-*gypsy* retrotransposons may have increased their abundance within the CWRs.

We found that most TE families are distributed across all analyzed beet species, indicating that the present beet repeat set can be traced back to the last common ancestor of both genera *Beta* and *Patellifolia*. After speciation from this ancestor, a few new TE families emerged, though rather sparsely. Instead, specific TE amplification/elimination has led to genus-and section-specific TE abundances. This global trend in beet genomes is mirrored in the ups and downs of individual TE families/lineages (e.g. SIRE/Maximus elements: Weber *et al*., 2010; chromoviruses: Weber *et al*., 2013; LINEs: Heitkam *et al*., 2014; non-autonomous LTR retrotransposons: Maiwald *et al*., 2021). Differences in the repeat profiles support the relationships of the current beet phylogenies (Frese and Ford-Lloyd, 2020; Sielemann *et al*., 2022) with characteristic repeatomes for species from the sections *Beta*, *Corollinae/Nanae*, and from the genus *Patellifolia*, respectively. However, whereas the amplification of TE families/lineages (in particular Ogre and non-autonomous LTR retrotransposons) has probably led to an increase in genome size of *Corollinae/Nanae* species, accumulation of specific TEs in the genomes of *Patellifolia* or section *Beta* species seem to have taken place at a comparable level, so that no striking differences in genome size are apparent between those beet taxa. To be more precise, there has been a strong amplification of few repeats in *Patellifolia* genomes, while many repeats have been amplified rather moderately in genomes of the section *Beta*.

Since the genomic shock of polyploidisation may stimulate rapid and dynamic genomic changes such as the activation of retrotransposons and viral elements (McClintock, 1984; Lopez-Gomollon *et al*., 2021), genome size may subsequently increase. The overall repeat content is indeed greater within the polyploid beet genomes compared to their closest diploid relatives, mainly due to an increased abundance of Ty3-*gypsy* retrotransposons. For most polyploid beet species, we measured larger genome sizes than mathematically expected, which can thus be attributed to a general accumulation of all kinds of present Ty3-*gypsy* retrotransposons instead of the targeted amplification of distinct TEs. During the ‘cycle of polyploidy’, polyploid plants usually undergo genome downsizing (Wendel, 2015), which may be the case for *B. macrocarpa* (since its genome size is smaller than mathematically expected) but not for the other polyploid beet species, indicating that these are relatively young polyploids (less than 0.9-1.4 million years; Romeiras *et al*., 2016).

It is often observed that genomic repeat profiles contain a phylogenetic signal (Dodsworth *et al*., 2015; McCann *et al*., 2020; Vitales *et al*., 2020b; Herklotz *et al*., 2021). In the case of *Beta* and *Patellifolia* species, this is only partly true: Overall, the repeatomes enable to separate the beet species into genera and sections. However, repeatome differences at the subspecies level do not reflect the currently proposed relationships of the subspecies taxa: As an example, *B. vulgaris* is considered to be more closely related to *B. maritima* accessions, regardless of their origin, than to *B. adanensis* (Wascher *et al*., 2022). According to kmer-based genomic distance, *B. adanensis* may even be considered a distinct species rather than a subspecies (Wascher *et al*., 2022). In contradiction with this report, focusing on the three subspecies *B. adanensis*, *B. vulgaris*, and *B. maritima*, the repeat profiles of the first two are most similar to each other. This may be explained by the *B. maritima* accessions used here: both Bmar1 and Bmar2 are from the Atlantic coast and are therefore less closely related to *B. vulgaris* than its presumed progenitor *B. maritima* from the Mediterranean area. From this, we conclude that the repeat profiles are not suited to address still debated phylogenetic questions regarding the (sub-)species level of beets. This also concerns the non-uniform treatment of *P. procumbens* and *P. webbiana* as distinct or as the same species.

The abundance of a distinct repeat depends on several factors that either cause an increase, a reduction, or that keep the current copy number stable. Those factors include an enhanced amplification (as mentioned above), the targeted or non-targeted elimination from the genome (e.g. by defense mechanisms and/or the loss of whole chromosomes or chromosome fragments), and selective pressure for the maintenance of specific genomic regions. An example in which some of these factors come into play is the section-specific abundance of endogenous pararetroviruses (EPRVs) found among the beets. There are three to five times more EPRVs within the genomes of wild beets from the section *Corollinae* compared to beets from the sister genus *Patellifolia*, the section *Beta*, and even *B. nana*. Such differences were also found between wild and cultivated potato species (Gaiero *et al*., 2019) and it was assumed that the EPRVs underwent either an increased amplification in wild potatoes or a selective bias in cultivated potatoes. As for sugar beet, we know that there is a trade-off between the targeted inactivation and a simultaneous preservation of EPRV sequences (Schmidt *et al*., 2021). Therefore, we believe that *Corollinae* species accumulated pararetroviral sequences by the gain of further EPRVs in comparison to the remaining beet species. Similar section-specific emergence and loss occurred also for other TE families/lineages, especially among the LTR retrotransposons.

### SatDNAs emerge, amplify and vanish without impacting the global genome size

The only repeat type for which no correlation with the beet genome size was found are the satDNAs. Strikingly, of all repeats, the satDNAs show the most pronounced specificity in abundance and distribution among the beet genomes, pointing to the fact that satDNAs are the most dynamic repeats (Garrido-Ramos, 2021). With this study, we present the first all-encompassing account of every present *Beta/Patellifolia* satDNA, involving all known as well as so far unpublished satDNAs. By comparison to publicly available databases, we determined that, with the exception of beetSat02-pEV (also found in quinoa: Schmidt *et al*., 2014), beetSat06-pAv34 (pRs34; also found in spinach: Dechyeva and Schmidt, 2006), and beetSat13 (ChenSat-1a; also found in quinoa: Heitkam *et al*., 2020), the satDNAs are specific for and restricted to beet species from the genera *Beta* and *Patellifolia*.

Usually, the repeatome of plants is constituted mostly by LTR retrotransposons, accounting for up to 80% of the plant genome size (Orozco-Arias *et al*., 2019). Only few plants are known in which the proportion of satDNAs is comparable to that of TEs (e.g. olive: Barghini *et al*., 2014; radish: He *et al*., 2015; *Fritillaria affinis*: Kelly *et al*., 2015). Here, we present sugar beet (*B. vulgaris*) as another plant species with peculiar high amounts of satDNAs. However, with diminishing phylogenetic relationship to sugar beet, the proportion of satDNAs decreases in its CWRs. In addition, each beet section/genus has its own distinct repeat profile, which sets it apart from the other (Fig. 5). Thus, we observed satDNAs specific for the genus *Patellifolia*, as well as for the sections *Corollinae/Nanae* (three different satDNAs, each) and *Beta* (one satDNA). Even the sole two satDNAs that are present in all analyzed beet species (i.e. beetSat02-pEV, beetSat06-pAV34) show variability in abundance as well as sequence and monomer length, resulting in genus- and section-specific variants that may initiate homogenization processes and sequence shifts in the future.

**Fig. 5:**
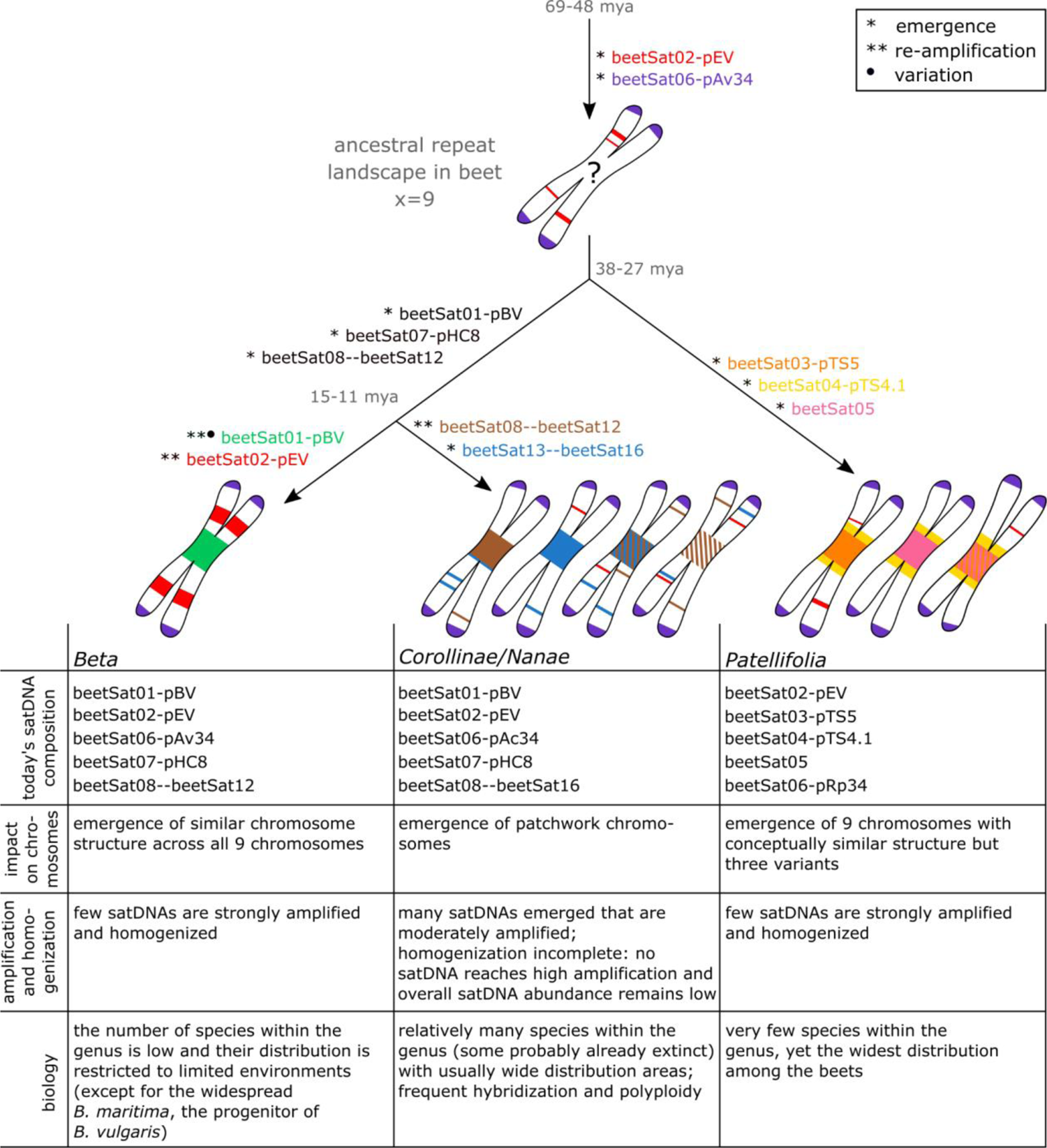
Scenario for the evolution of satDNAs during the history of beet speciation. As the satDNAs beetSat02-pEV and beetSat06 are also present in the spinach/quinoa outgroup (Dechyeva and Schmidt, 2006; Schmidt *et al*., 2014), we assume an emergence of these satDNAs even before the split of the beet crown. All other identified satDNAs seem to have appeared earliest in the direct beet ancestor with the subsequent emergence of genus- and section-specific satDNAs and different amplification and differentiation patterns. The age of the respective beet taxa has been estimated in million years ago (mya) by Hohmann *et al*. (2006).

As beetSat02-pEV and beetSat06-pAV34 are distributed among all analyzed beets (Fig. 5) and as variants of them are also present in more distantly related plant species (beetSat02-pEV in quinoa: Schmidt *et al*., 2014; beetSat06-pAV34 in spinach: Dechyeva and Schmidt, 2006), we assume that these satDNAs were already present in the common ancestor of both genera *Beta* and *Patellifolia*. During the subsequent beet speciation, a great variety of new satDNAs emerged and some of them accumulated section-specifically by remarkable re-amplification. Such a restriction to individual sections or species is sometimes known for TEs, but is particularly pronounced for the fast evolving satDNAs (Orozco-Arias *et al*., 2019, Garrido-Ramos, 2021). For example, beetSat13’s specificity in abundance is visible even at the level of beet accessions (Bmrh1 vs. Bmrh2). Moreover, this satDNA can be found in all *Corollinae/Nanae* species as well as in the more distantly related quinoa (ChenSat-1a: Heitkam *et al*., 2020), but not in beet genomes from the genus *Patellifolia* and the section *Beta* (Fig. 5). The patchy distribution of beetSat13 among Amaranthaceae members may be either explained by its loss in most of the beet genomes except those from the section *Corollinae/Nanae*, or by an independent acquisition into the genome of the *Corollinae/Nanae* ancestor. It was argued that in quinoa this satDNA emerged from a CACTA-like TE (Belyayev *et al*., 2020). Thus, an independent acquisition/generation of beetSat13 into/within the genome of the *Corollinae/Nanae* ancestor may have occurred similarly by spawning from a TE or even by the transmission of extrachromosomal circular molecules consisting of this satDNA (Navrátilová *et al*., 2008).

### SatDNAs impact beet chromosome architecture in a gene pool-dependent manner

Apart from their role as intercalary heterochromatin (beetSat02-pEV) and (sub-)telomere contributors (beetSat06-pAV34), satDNAs often provide the DNA backbone onto which the centromeres are formed (Melters *et al*., 2013; Garrido-Ramos, 2021). Strikingly, this iconic location is not occupied by the same satDNA among beets, but by a peculiarly high number of different satDNAs (Fig. 5). It is not unusual for the centromeric satDNAs to differ even in very closely related species (Henikoff *et al*., 2001; Melters *et al*., 2013). However, whereas the centromeric satDNA in beets from the section *Beta* is well defined and conserved across the whole karyotype (Schmidt and Metzlaff, 1991; Zakrzewski *et al*., 2011), beets from the section *Corollinae/Nanae* and the genus *Patellifolia* seem to differ in their satDNA composition even from chromosome to chromosome. This phenomenon was previously observed in potatoes (Gong *et al*., 2012) and especially in legumes (Avila Robledillo *et al*., 2020), as well as in animal species (chicken: Shang *et al*., 2010). Usually, the most abundant satDNA of a species is the one constituting the centromeres (Melters *et al*., 2013). Yet, while this is true for beets from the section *Beta*, there is no clearly predominant satDNA in beets from the section *Corollinae/Nanae* and the genus *Patellifolia*, going along with the observation that the variety of similarly abundant satDNAs is spread over the (peri-)centromeric regions of subsets of chromosomes. This is particularly remarkable as these satDNAs (i.e. beetSat07-pHC8 to beetSat11, beetSat13, beetSat15, and beetSat17 in *Corollinae/Nanae* species; beetSat03-pTS5 and beetSat05 in *Patellifolia* species) differ considerably in their monomer lengths and/or sequences and yet chromosomes maintain an error-free cell division. Furthermore, centromeric satDNA families of other closely related plant taxa (e.g. *Arabidopsis* species), albeit species-specific, nevertheless derived from a common ancestor (summarized by Garrido-Ramos, 2021), which does not seem to be true for the beets: Monomer sequence and length differences are too large to be explained by high mutation rates, even though some satDNAs (e.g. beetSat01-pBV) indeed evolve towards quite considerable changes in monomer sequence and length (see Additional file 1: Fig. S4). The centromere drive model (Henikoff *et al*., 2001) does not seem to apply either, as the presence of multiple centromeric satDNAs with different sequences precludes any sequence-dependent coevolution with the kinetochore complex (Avila Robledillo *et al*., 2020). Considering the evolutionary mode of satDNAs, which are thought to typically evolve in a concerted manner (Šatović-Vukšić and Plohl, 2023), it may be that *Corollinae/Nanae* and *Patellifolia* species are only at an intermediate stage on the way to centromere and satDNA homogenization. This is supported by the fact that beet species from the section *Corollinae/Nanae* in particular belong to a highly variable hybrid complex (Frese and Ford-Lloyd, 2020). Hence, hybridization events between genotypes with different satDNA profiles may frequently disrupt any ongoing satDNA homogenization processes. This may have led to the observed variety of lowly amplified satDNAs, which were then selectively retained to form unique satDNA mixes at each centromere within one nucleus. This would then result in the observed combination of different chromosome sets with different satDNA architectures. To account for their unique chromosome make-up, we refer to the *Corollinae/Nanae* chromosomes as patchwork chromosomes (Fig. 5; middle). Alternatively, keeping a set of different centromeres may hold an advantage for these beets, especially in chromosome recognition during meiotic pairing. For the *Corollinae/Nanae*, we cannot yet determine if there is chromosome-complementarity among individual satDNAs. Nevertheless, multicolor FISH indicates a possible chromosome-complementarity of beetSat03-pTS and beetSat05 in the genus *Patellifolia*. Taking together, we detect vastly different satDNA landscapes that vary in satDNA family number, abundance and location. These provide the DNA backbone for the formation of overall different chromosome architectures within a nucleus, leading from overall similar chromosome/centromere structures (section *Beta*), over conceptually similar chromosomes (*Patellifolia*) towards completely different chromosomal/centromeric setups (*Corollinae/Nanae*).

## Conclusion

In the use of wild germplasm for breeding, crossing barriers often impede the sustainable introgression of target traits from wild species into crops. This is also the case for cultivated and wild beet species, which are taxa with isokaryotypic chromosomes, but which nevertheless exhibit strong crossing borders between gene pools. We found that global differences between beet genomes can be attributed primarily to their repeatomes, especially to the specific composition of satDNAs, whereas the base number and morphology of chromosomes is rather consistent. After the divergence from the common beet ancestor, genus- and section-specific repeat variants propagated by independent transposition and centromeres evolved distinctly, corresponding to the beet phylogeny and the categorization into gene pools. Based on their repeatome, beet species divide into three different groups (section *Beta* as primary gene pool, section *Corollinae/Nanae* as secondary gene pool, and genus *Patellifolia* as tertiary gene pool), supporting the repeated handling of *B. nana* as a member of the section *Corollinae*. In comparison to the other beet species, this section is characterized by a peculiarly high number of different satDNAs with comparable abundances. We found that these satDNAs contribute to the *Corollinae/Nanae* centromeres in a composite manner, resulting in a highly variable chromosome composition (patchwork chromosomes). These insights are valuable for future beet breeding as they help to face the challenge of overcoming postzygotic crossing barriers and provide unique insights into genome, repeatome, and chromosome evolution in karyotypically stable taxa.

## METHODS

### Plant material, genome size measurement, and DNA extraction

The 17 investigated beet accessions cover the whole spectrum of species within the genera *Beta* and *Patellifolia* (see Table 1). Seeds of the *B. vulgaris* subsp. *vulgaris* genotype KWS2320 were obtained from KWS Saat SE, Einbeck, Germany. The seeds of all other accessions were obtained from the Leibniz Institute of Plant Genetics and Crop Plant Research Gatersleben, Germany. The material of the KWS SAAT SE & Co. KGaA, Einbeck and IPK Gatersleben was transferred under the regulations of the standard material transfer agreement (SMTA) of the International Treaty. The plants were grown under long day conditions in a greenhouse.

Nuclei extraction and staining were performed using the CyStain PI OxProtect reagent kit from Sysmex. Nuclear DNA was measured by flow cytometry, using *Raphanus sativus* (2C = 1.11 pg DNA) or *Pisum sativum* (2C = 9.07 pg DNA; Doležel *et al*., 1992) as internal standard. Four DNA estimations were carried out for each plant (5000 nuclei per analysis) on at least two different days. Nuclear DNA content (2C value in [pg]) was calculated as: sample peak mean/standard peak mean × 2C DNA content of the standard. DNA amounts in picograms were converted to the number of base pairs using the conversion factor 1 pg DNA = 0.978 × 109 bp (Doležel *et al*., 2003). The mean nuclear DNA content was then calculated for each plant as 1C value (in [Mbp]; Table 1).

Genomic DNA was extracted from approximately 100 mg of fresh leaf tissue samples using the NucleoSpin® Plant II protocols from Macherey-Nagel. Libraries were prepared using the TruSeq Nano DNA library preparation kit (Fa. Illumina) and were sequenced on an Illumina HiSeq1500 sequencer at the CeBiTec Sequencing Core Facility, Bielefeld University, Germany (250 bp paired-end reads).

### Repeat classification and quantification

The Illumina reads were trimmed to a length of 100 bp using Trimmomatic (v0.39; Bolger *et al*., 2014). High quality of the trimmed reads was ensured by FastQC examination (v0.11.5; Andrews, 2020). Bowtie2 (v2.2.6; Langmead and Salzberg, 2012) was applied to identify and subsequently remove reads representing organellar DNA from the sequence data by mapping them to a database containing publicly available chloroplast and mitochondrial DNAs of *Beta* species (https://ncbi.nlm.nih.gov/nuccore). The sequence reads were then subsampled to reduce the genome coverage to 0.1× for all species, and different numbers of paired-end reads sampled depending on the genome size. To standardize all read pre-treatments, we have used the mentioned tools embedded in the preparation module of the ECCsplorer pipeline (Mann *et al*., 2022). For the identification and quantification of repetitive sequences, the similarity-based read clustering method was applied as described by Novák *et al*. (2010) and implemented in the RepeatExplorer2 pipeline (Novák *et al*., 2013, 2020). We used the pipeline default settings (90% similarity over 55% of the read length) and included our custom database of Betoideae repeats (https://zenodo.org/record/8255813) for individual analyses (each beet accession distinctly) as well as an all-vs-all comparison across all species (accession 1 of every species; see Table 1). When comparing different accessions of the same species (Bmar1/Bmar2, Bmrh1/Bmrh2, Bint1/Bint2, and Bnan1/Bnan2), the RepeatExplorer2 pipeline produced highly similar results for the respective repeat compositions (see Additional file 1: Table S1). Any discrepancies resulted from the read sampling as we confirmed by using re-sampled read datasets as input. Hence, only one accession per species was included in the comparative RepeatExplorer2 analysis.

For this all-vs-all comparison, each read set was downsampled to represent 4% of the respective genome (coverage of 0.04×) based on ‘ploidy corrected genome sizes’ (1C values [Table 1] divided by 2 [for the tetraploid species *B. macrocarpa*, *B. corolliflora*, and *P. patellaris*] and 2.5 [for the pentaploid species *B. intermedia*], respectively). Nevertheless, the pipeline reached a limit, retrieving 3,985,090 reads. Although automatic annotation of read clusters was improved by the inclusion of our custom Betoideae repeat database, not all clusters were classified. The cluster graph shapes of all unassigned clusters that represented at least 0.001% of the investigated genomes, were examined manually and consensus sequences, if provided, were searched using publicly available databases (https://blast.ncbi.nlm.nih.gov/Blast.cgi). Finally, the relative abundance of each repeat was calculated based on the number of reads in the respective cluster, excluding remaining organellar reads.

### Sequence comparison

Multiple sequence alignments were generated with the MAFFT (v7.017; Katoh and Standley, 2013) and MUSCLE (v3.8.31; Edgar, 2004) local alignment tools. They have been manually refined and used for the calculation of pairwise sequence identities. We explored and visualized sequences with the multipurpose software Geneious 6.1.8 (Kearse *et al*., 2012). Dotplots were generated with FlexiDot (Seibt *et al*., 2018) with a word size of 20 bp using long reads of *B. vulgaris* (PacBio; accession number SRX3402137; Funk *et al*., 2018), *B. corolliflora* (ONT; accession number ERS13530775; Sielemann *et al*., 2023a), and *P. procumbens* (ONT; accession number ERS13530778; Sielemann *et al*., 2023a).

### Generation of satDNA probes

The probe for the *Patellifolia*-specific satDNA beetSat05 was ordered as EXTREmer oligonucleotides synthesized by *Eurofins Genomics* based on the RepeatExplorer2 consensus sequence (Additional file 1: Data S1).

Standard PCR reactions of genomic *B. corolliflora* (BETA 408) and *B. nana* (BETA 546) DNA were performed using primer pairs designed for two further beet-specific satDNAs (beetSat15 and beetSat17; Additional file 1: Table S3). The PCR conditions were 94 °C for 3 min followed by 35 cycles of 94 °C for 30 s, primer-specific annealing temperature for 30 s, 72 °C for 25 sec (beetSat15) and 45 s (beetSat17), respectively, and a final incubation at 72 °C for 5 min. PCR fragments were purified, cloned and commercially sequenced. Sequenced inserts spanning two (beetSat17; 279 bp) and six (betSat15; 303 bp) monomers of the respective satDNA were used as probes for the following hybridization experiments. Their identity to the respective reference sequence (RepeatExplorer2 consensus sequence) was 96.4% (beetSat15) and 93.7% (beetSat17), respectively.

### Preparation of chromosome spreads and fluorescent in situ hybridization (FISH)

The meristem of young leaves was used for the preparation of mitotic chromosomes. The plant tissues were treated as described by Schmidt *et al*. (2023). Accumulation of metaphases was achieved by a combination of an incubation with 2 mM 8-hydroxyquinoline for 3 h and nitrous oxide for 30 min. Fixed plant material was digested using the ‘leaf enzyme solution II’ (Schmidt *et al*., 2023).

The probes for the chromosomal localization of several satDNAs using FISH were labeled directly as well as indirectly: The probes ‘pBV I’ (Schmidt and Metzlaff, 1991; Kubis *et al*., 1998) and ‘beetSat05’ (this manuscript; Additional file 1: Data S1) for specific centromeric satDNA families were directly labeled with DY647-dUTP (Dyomics). The probe ‘pEV I’ (similar to Schmidt *et al*., 1991; accession number OY726583) marking an intercalary satDNA family (Kubis *et al*., 1998) was directly labeled with DY415-dUTP (Dyomics). The probe ‘pTS4.1’ for the pericentromeric satDNA in *Patellifolia* species (Schmidt *et al*., 1990; Schmidt and Heslop-Harrison, 1996) was indirectly labeled by PCR in the presence of digoxigenin-11-dUTP detected by antidigoxigenin-fluorescein isothiocyanate (FITC; both from Roche Diagnostics). The remaining satDNAs (‘pTS5’: Schmidt *et al*., 1990, Schmidt and Heslop-Harrison, 1996; ‘pHC8’: Gindullis *et al*., 2001; ‘pRN1’: Kubis *et al*., 1997; beetSat08: accession number OY726584 [corresponding to ‘BlSat1’ from Ha, 2018]; beetSat13 [corresponding to ‘ChenSat-1a’]: Heitkam *et al*., 2020; beetSat15 and beetSat17: this manuscript, accession numbers OY726585 and OY726586) were indirectly labeled by PCR in the presence of biotin-16-dUTP (Roche Diagnostics) detected by streptavidin-Cy3 (Sigma–Aldrich). The hybridization procedure was performed as described previously (Schmidt *et al*., 1994) with a stringency of 82%. Chromosomes were counterstained with DAPI (4’,6’-diamidino-2-phenylindole; Böhringer, Mannheim) and mounted in an antifade solution (CitiFluor). Slides were examined with a fluorescence microscope (Zeiss Axioplan 2 imaging) equipped with appropriate filters. Images were acquired directly with the Applied Spectral Imaging v. 3.3 software coupled to the high-resolution CCD camera ASI BV300-20A. After separate capture for each fluorochrome, the individual images were combined computationally and processed using Adobe Photoshop CS5 software (Adobe Systems, San Jose, CA, USA). We used only contrast optimization, Gaussian and channel overlay functions affecting all pixels of the image equally.

## DECLARATIONS

### Ethics approval and consent to participate

The material of the KWS SAAT SE & Co. KGaA, Einbeck and IPK Gatersleben was transferred under the regulations of the standard material transfer agreement (SMTA) of the International Treaty. Plants were grown in accordance with German legislation.

### Consent for publication

Not applicable.

### Availability of data and materials

Illumina whole genome sequence data are available at EBI under accession numbers as indicated in Table 1. ONT reads for *B. corolliflora* and *P. procumbens* are available at EBI under the accession numbers ERS13530775 and ERS13530778. RepeatExplorer2 outputs have been made available at ZENODO (doi: 10.5281/zenodo.7821055; https://zenodo.org/record/7821055). Satellite DNA consensus sequences and the sequences used as FISH probes are available in Additional file 1: Data S1 and Data S2. Furthermore, the cloned sequences of the satDNAs beetSat02-pEV and beetSat08, as well as the newly identified satDNAs beetSat15 and beetSat17 were submitted to ENA (accession numbers OY726583-OY726586).

### Competing interests

The authors declare no competing interests.

### Funding

This work was supported by the German Federal Ministry of Education and Research (call „Epigenetics: Opportunities for Plant Research“, grant 031B1221A). KS was funded by Bielefeld University.

### Authors’ contributions

NS, BW, DH, and TH designed the study. NS selected and cultivated the plants. NS, KS, and BP performed DNA extraction. NS and JF performed flow cytometry. BP performed sequencing. NS and SB performed FISH. NS implemented the bioinformatic methodology, analyzed the data and prepared the figures and tables. NS, KS, and TH wrote the manuscript. All authors read and approved the final manuscript.

## Acknowledgements

We acknowledge the KWS SAAT SE & Co. KGaA, Einbeck and the Genbank Gatersleben for providing seeds and data for the investigated accessions. Computational resources for RepeatExplorer2 analysis were provided by the ELIXIR-CZ project (LM2015047), part of the international ELIXIR infrastructure.

## SUPPLEMENTARY DATA

Additional supporting information may be found in the online version of this article and consists of the following. Fig. S1: DAPI-stained chromosomes of 17 different beet accessions. Fig. S2: Correlation between repeat proportion and genome size in *Beta* and *Patellifolia* species. Fig. S3: Comparative repeat composition among *Beta* and *Patellifolia* species. Fig. S4: Arrangement of beet satDNA monomers on long reads of *B. vulgaris* (Bvul), *B. corolliflora* (Bcor), and *P. procumbens* (Ppro). Fig. S5: Comparative nucleotide alignment of the three section-specific beetSat02-pEV variants. Table S1: Genome proportion [%] of different repeat classes and superfamilies (repeat proportion) in *Beta* and *Patellifolia* species. Table S2: Genome proportion [%] of different satDNAs (and tandem repeats) across *Beta* and *Patellifolia* species. Table S3: Primer sequences for the amplification of beet-specific satellite DNAs. Data S1: beetSat nucleotide sequences derived as RepeatExplorer2 consensuses from the comparative analysis. Data S2: beetSat sequences used as FISH probes.

## Notes

https://zenodo.org/record/7821055

### Competing Interest Statement

The authors have declared no competing interest.

### Summary of Updates

We unfortunately had a typo in one of the author's names in the PDF version of the manuscript.

