## Supplementary Data for "Repeat turnover meets stable chromosomes: repetitive DNA sequences mark speciation and gene pool boundaries in sugar beet and wild beets"

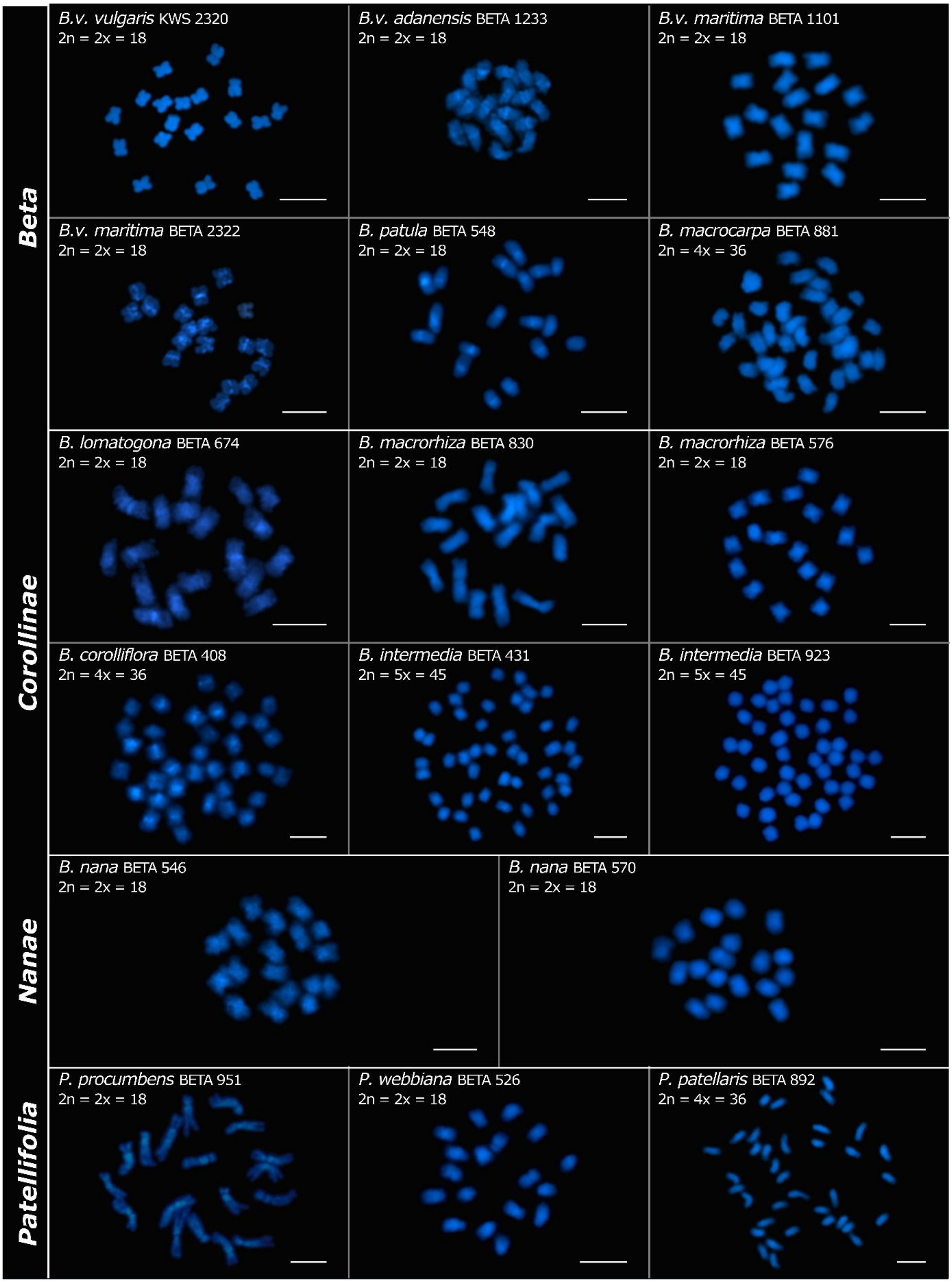


**Fig. S1:** DAPI-stained chromosomes of 17 different beet accessions. The diploid chromosome set comprises 18 chromosomes. Tetraploid beet species have 36 chromosomes and the pentaploid *B. intermedia* has 45 chromosomes. Scale bars = 5 µm.

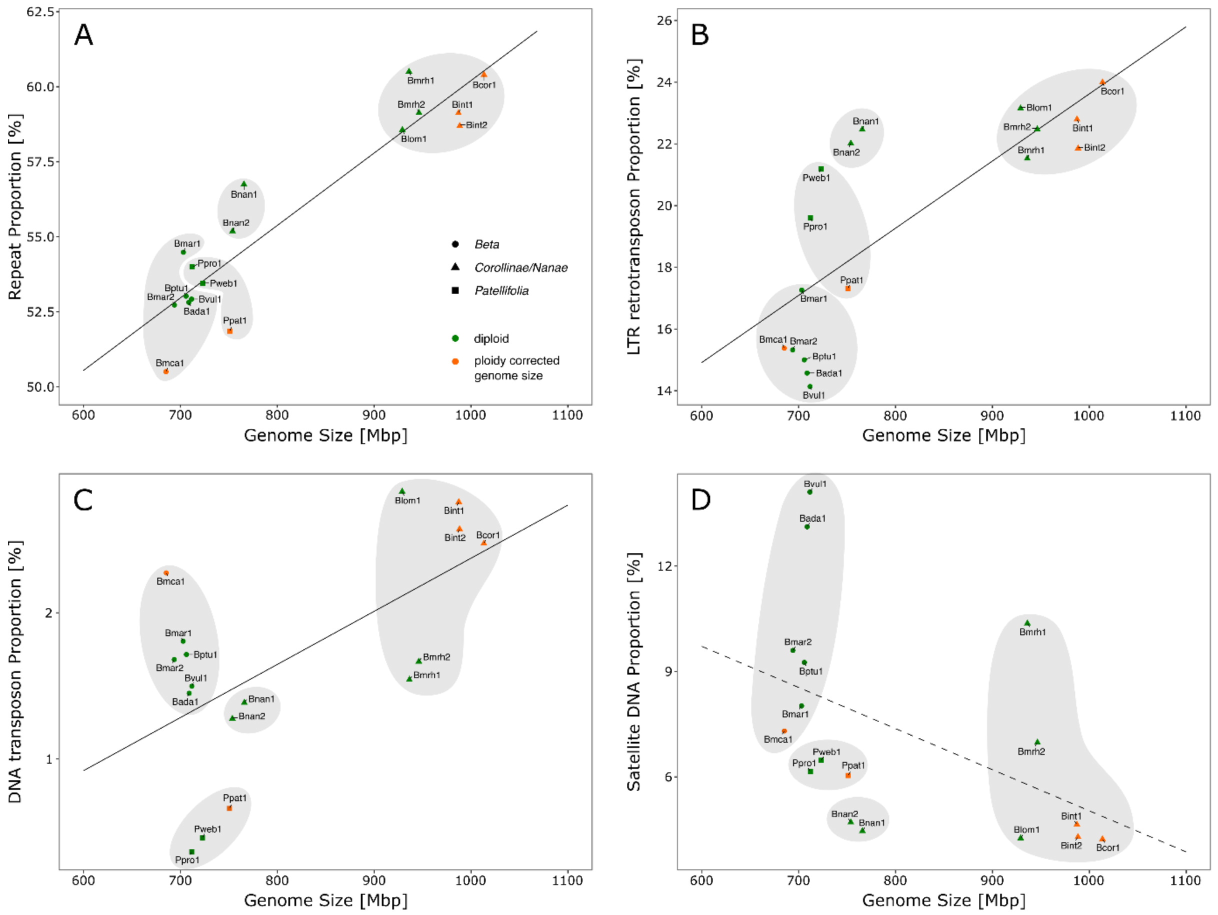


**Fig. S2:** Correlation between repeat proportion and genome size in *Beta* and *Patellifolia* species. For better comparability with natural diploids (green), polyploid species were downsampled as artificial diploids (orange). Shapes and shades indicate the different beet sections and genera, respectively. The sample abbreviations are according to Table 1. (A) Positive correlation between the total repeat proportion and genome size with $r^2 = 0.927$ ($p < 9.26e^{-8}$). (B) Positive correlation between LTR retrotransposon proportion and genome size with $r^2 = 0.773$ ($p = 2.77e^{-4}$). (C) Positive correlation between DNA transposon proportion and genome size with $r^2 = 0.609$ ($p < 9.41e^{-3}$). (D) The satellite DNA proportion and genome size tend to relate negatively with $r^2 = -0.474$ ($p > 0.05$).

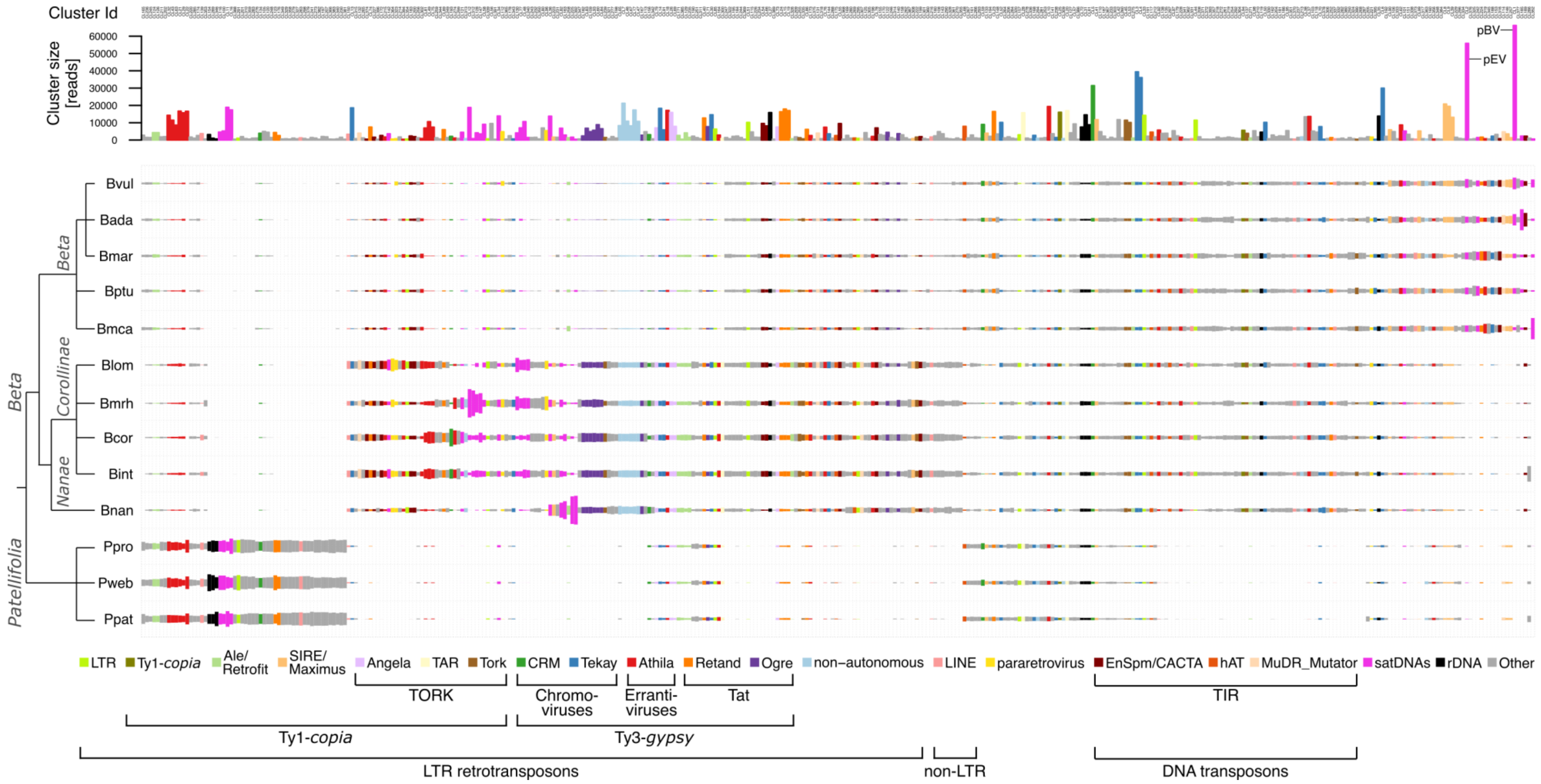


**Fig. S3:** Comparative repeat composition among *Beta* and *Patellifolia* species. The bars represent the distribution of clusters comprising at least 400 reads ( ≥ 0.01% of the analyzed reads) among the analyzed Betoideae members. Rectangles are colored according to the type of repetitive element and their size is proportional to the genomic abundance in the respective species (abbreviations as in Table 1; accession 1 of the respective species). The satellite DNAs beetSat01-pBV and beetSat02-pEV are indicated as clusters comprising the highest amount of reads.

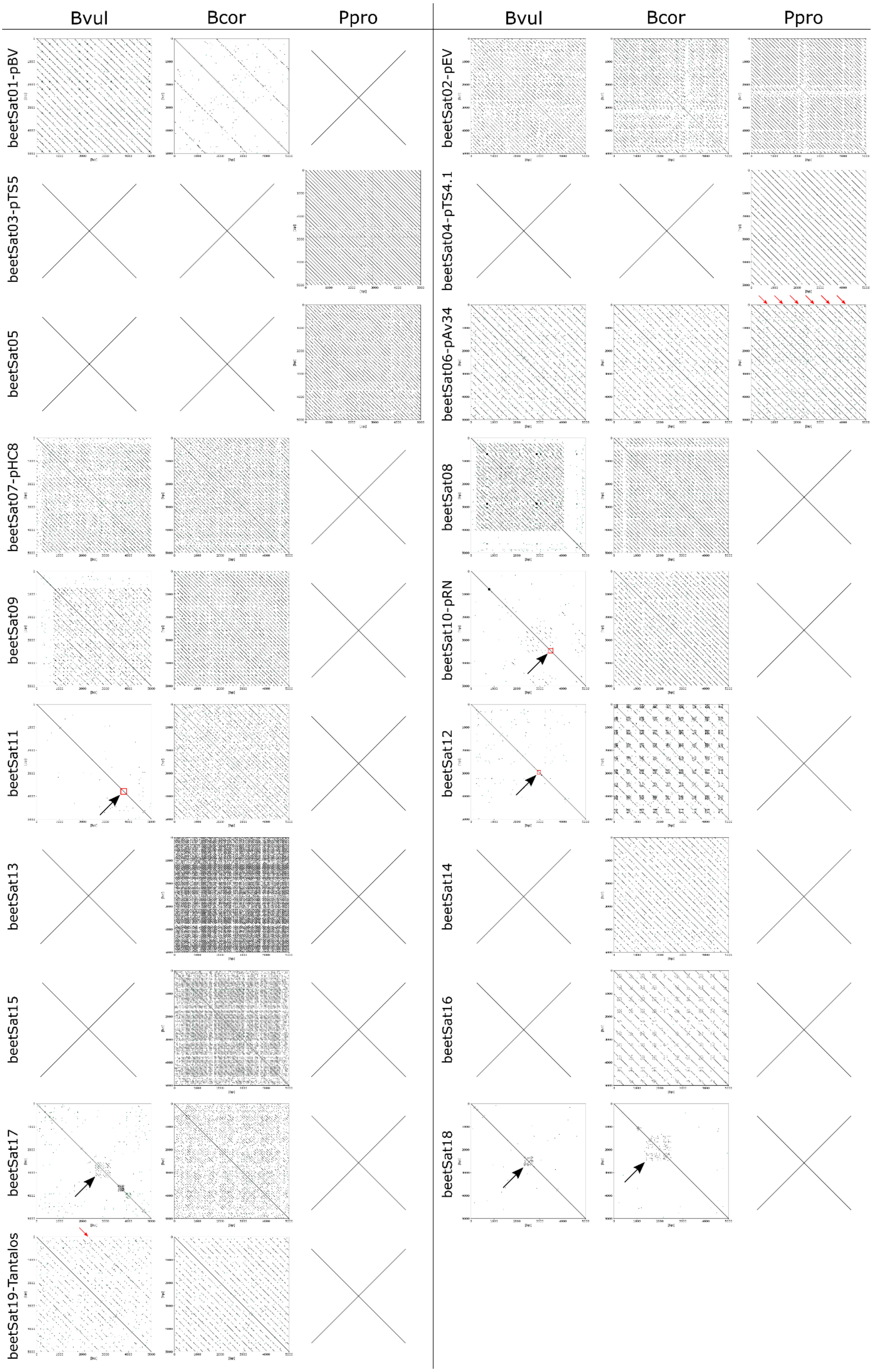
Bvul
Bcor
Ppro
Bvul
Bcor
Ppro
beetSat01-pBV
beetSat02-pEV
beetSat03-pTS5
beetSat04-pTS4.1
beetSat05
beetSat06-pAv34
beetSat07-pHC8
beetSat08
beetSat09
beetSat10-pRN
beetSat11
beetSat12
beetSat13
beetSat14
beetSat15
beetSat16
beetSat17
beetSat18
beetSat19-Tantalos

**Fig. S4**: Arrangement of beet satDNA monomers on long reads of *B. vulgaris* (Bvul; PacBio; accession number SRX3402137), *B. corolliflora* (Bcor; ONT; accession number ERS13530775), and *P. procumbens* (Ppro; ONT; accession number ERS13530778). A field was crossed out if no hits were found in the respective read set. The monomers are arranged in continuous arrays, short arrays (beetSat08, beetSat09, beetSat17, and beetSat18 in Bvul, as well as beetSat18 in Bcor), higher order arrangements (red arrows), or as non repetitive sequences (red rectangles). The satDNA beetSat18 occurs only in the genome of *B. nana* in a tandemly arranged manner which was proven using sequenced reads from *B. nana* (data not shown). If detected, we show a self dotplot of a representative excerpt over a 5000 bp window. Forward matches are marked by black lines, whereas reverse matches are indicated in green. To account for the ONT read error rate of 5-10%, we chose a word size of 20 bp and allowed five mismatches.

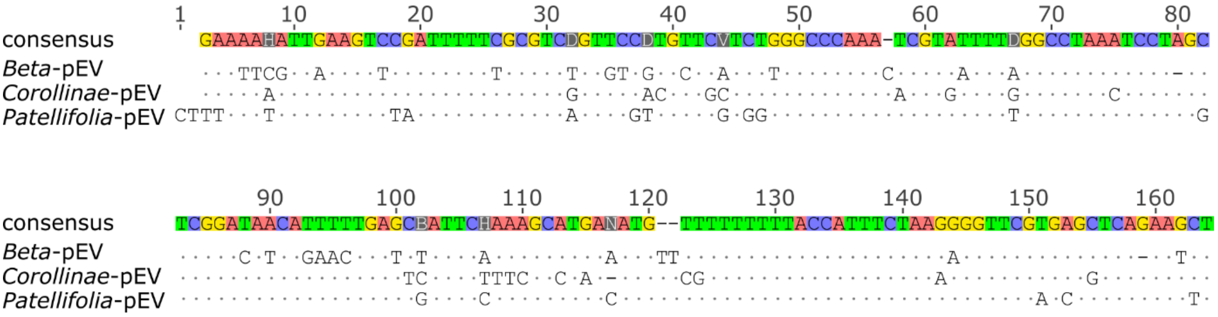


**Fig. S5:** Comparative nucleotide alignment of the three section-specific beetSat02-pEV variants. Deviations from the consensus sequence are indicated. The *Beta*-specific beetSat02-pEV sequence corresponds to the 160 bp long monomer of the EcoRI satDNA described by Schmidt *et al.* (1991; accession number Z22848.1), and the *Patellifolia*-specific beetSat02-pEV sequence corresponds to the 161 bp long monomer of pAp11-1 described by Dechyeva *et al.* (2003; accession number AJ414554.1). The *Corollinae*-specific beetSat02-pEV sequence was retrieved during the comparative cluster analysis by the RepeatExplorer2 software: Based on a cluster annotated as the satDNA beetSat02-pEV, exclusively comprising reads from Blom, Bmrh, Bcor, and Bint, the monomer sequence was derived as a consensus sequence from the read assembly.

**Table S1:** Genome proportion [%] of different repeat classes and superfamilies (repeat proportion) in *Beta* and *Patellifolia* species. Abbreviations (species) as in Table 1. [1] Repeat proportion based on real ploidy. [2] More details in Table S2.

| species | ploidy | genome size based on corrected ploidy | total repeat proportion[1] | total repeat proportion based on corrected ploidy | total proportion of LTR retrotransposons[1] | total proportion of LTR retrotr. based on corrected ploidy | proportion of Ty1-*copia* retrotransposons[1] | | | | | | | | proportion of Ty3-*gypsy* retrotransposons[1] | | | | | | proportion of non-autonomous LTR retrotransposons[1] |
|---|---|---|---|---|---|---|---|---|---|---|---|---|---|---|---|---|---|---|---|---|---|
| | | | | | | | total proportion of Ty1-*copia* retrotransposons | Ale/Retrofit | Bianca | Ivana/Oryco | SIRE/Maximus | Angela | TAR | Tork | total proportion of Ty3-*gypsy* retrotransposons | CRM | Tekay | Athila | Retand | Ogre | |
| Bvul1 | 2n=2x | - | 52.92 | - | **14.13** | - | 6.44 | 0.24 | 0.26 | 0.00 | 4.13 | 0.32 | 0.68 | 0.37 | 6.02 | 1.07 | 2.46 | 1.41 | 0.67 | 0.00 | 0.05 |
| Bada1 | 2n=2x | - | 52.81 | - | **14.57** | - | 6.60 | 0.27 | 0.20 | 0.41 | 4.33 | 0.32 | 0.33 | 0.43 | 7.33 | 1.25 | 2.73 | 1.97 | 1.01 | 0.00 | 0.10 |
| Bmar1 | 2n=2x | - | 54.48 | - | **17.27** | - | 6.28 | 0.33 | 0.26 | 0.00 | 3.01 | 0.40 | 0.62 | 0.75 | 8.89 | 0.22 | 2.73 | 2.18 | 2.64 | 0.00 | 0.46 |
| Bmar2 | 2n=2x | - | 52.73 | - | **15.32** | - | 6.25 | 0.27 | 0.22 | 0.00 | 3.25 | 0.64 | 0.57 | 0.52 | 7.62 | 0.36 | 2.82 | 2.42 | 1.75 | 0.00 | 0.28 |
| Bptu1 | 2n=2x | - | 53.02 | - | **15.00** | - | 5.70 | 0.27 | 0.29 | 0.30 | 3.30 | 0.48 | 0.49 | 0.40 | 7.82 | 0.61 | 2.69 | 2.07 | 1.84 | 0.00 | 0.36 |
| Bmca1 | 2n=4x | 685 Mbp | 57.99 | 50.51 | **19.71** | 15.37 | 7.27 | 0.39 | 0.38 | 0.03 | 3.57 | 0.49 | 0.64 | 1.18 | 10.26 | 0.84 | 3.70 | 2.48 | 2.24 | 0.00 | 0.11 |
| Blom1 | 2n=2x | - | 58.56 | - | **23.15** | - | 4.90 | 0.11 | 0.17 | 0.13 | 1.04 | 1.42 | 0.84 | 0.50 | 15.17 | 0.35 | 5.27 | 4.92 | 2.67 | 1.69 | 3.08 |
| Bmrh1 | 2n=2x | - | 60.51 | - | **21.52** | - | 4.76 | 0.39 | 0.20 | 0.09 | 1.42 | 1.16 | 0.48 | 0.28 | 12.61 | 0.43 | 3.12 | 4.05 | 1.90 | 2.75 | 3.66 |
| Bmrh2 | 2n=2x | - | 59.13 | - | **22.48** | - | 5.32 | 0.28 | 0.22 | 0.08 | 1.63 | 1.18 | 0.56 | 0.26 | 12.70 | 0.41 | 3.22 | 4.07 | 1.65 | 2.77 | 3.76 |
| Bcor1 | 2n=4x | 1014 Mbp | 67.65 | 60.40 | **30.06** | 23.99 | 5.99 | 0.67 | 0.26 | 0.08 | 1.20 | 1.26 | 0.95 | 0.78 | 17.60 | 0.65 | 3.70 | 5.78 | 3.42 | 3.43 | 4.80 |
| Bint1 | 2n=5x | 987 Mbp | 68.30 | 59.12 | **29.54** | 22.80 | 6.14 | 0.63 | 0.21 | 0.05 | 1.41 | 1.40 | 1.03 | 0.59 | 17.78 | 0.98 | 4.18 | 5.83 | 3.35 | 2.69 | 4.14 |
| Bint2 | 2n=5x | 988 Mbp | 68.49 | 58.70 | **29.23** | 21.85 | 6.03 | 0.58 | 0.23 | 0.48 | 1.58 | 1.35 | 0.99 | 0.49 | 16.96 | 1.21 | 3.82 | 5.35 | 2.89 | 3.31 | 4.18 |
| Bnan1 | 2n=2x | - | 56.75 | - | **22.46** | - | 4.76 | 0.36 | 0.23 | 0.00 | 1.06 | 1.21 | 0.48 | 0.52 | 12.57 | 0.61 | 4.84 | 2.02 | 1.50 | 3.04 | 4.90 |
| Bnan2 | 2n=2x | - | 55.19 | - | **22.01** | - | 5.23 | 0.26 | 0.21 | 0.00 | 1.84 | 1.33 | 0.42 | 0.45 | 12.28 | 0.47 | 5.03 | 2.36 | 1.04 | 2.98 | 4.43 |
| Ppro1 | 2n=2x | - | 54.01 | - | **19.60** | - | 4.17 | 0.54 | 0.15 | 0.00 | 1.15 | 0.74 | 1.27 | 0.00 | 14.37 | 1.53 | 1.68 | 8.49 | 2.53 | 0.12 | 0.03 |
| Pweb1 | 2n=2x | - | 53.45 | - | **21.19** | - | 4.31 | 0.65 | 0.18 | 0.00 | 1.09 | 0.72 | 1.31 | 0.04 | 15.62 | 1.53 | 1.85 | 8.96 | 3.07 | 0.15 | 0.05 |
| Ppat1 | 2n=4x | 751 Mbp | 59.02 | 51.86 | **22.37** | 17.31 | 4.96 | 0.85 | 0.33 | 0.00 | 1.30 | 0.74 | 1.23 | 0.12 | 17.20 | 1.46 | 3.64 | 7.85 | 3.25 | 0.27 | 0.05 |

**Table S1:** continued

| species | proportion of LINEs[1] | proportion of SINEs[1] | proportion of pararetroviruses[1] | proportion of DNA transposons: total proportion of DNA transposons[1] | total proportion of DNA transposons based on corrected ploidy | EnSpm/CACTA[1] | hAT[1] | MuDR_Mutator[1] | PIF/Harbinger[1] | Helitron[1] | total proportion of satellite DNAs[1,2] | total proportion of satellite DNAs based on corrected ploidy | total proportion of rDNA[1] |
|---|---|---|---|---|---|---|---|---|---|---|---|---|---|
| Bvul1 | 0.13 | 0.05 | 0.24 | **1.50** | - | 0.73 | 0.32 | 0.29 | 0.09 | 0.06 | 13.70 | - | 1.25 |
| Bada1 | 0.13 | 0.05 | 0.00 | **1.45** | - | 0.53 | 0.30 | 0.43 | 0.08 | 0.11 | 12.86 | - | 1.24 |
| Bmar1 | 0.13 | 0.10 | 0.11 | **1.81** | - | 0.93 | 0.46 | 0.26 | 0.11 | 0.05 | 7.57 | - | 2.60 |
| Bmar2 | 0.13 | 0.08 | 0.18 | **1.68** | - | 0.87 | 0.30 | 0.27 | 0.08 | 0.15 | 9.29 | - | 2.10 |
| Bptu1 | 0.16 | 0.06 | 0.14 | **1.72** | - | 0.88 | 0.30 | 0.36 | 0.12 | 0.06 | 8.80 | - | 2.31 |
| Bmca1 | 0.24 | 0.10 | 0.06 | **2.45** | 2.28 | 1.25 | 0.52 | 0.25 | 0.19 | 0.24 | 7.07 | 7.31 | 1.69 |
| Blom1 | 0.27 | 0.31 | 0.60 | **2.83** | - | 1.78 | 0.30 | 0.40 | 0.08 | 0.26 | 3.97 | - | 1.51 |
| Bmrh1 | 0.26 | 0.23 | 0.87 | **1.54** | - | 0.97 | 0.19 | 0.12 | 0.12 | 0.15 | 10.22 | - | 1.85 |
| Bmrh2 | 0.12 | 0.26 | 0.94 | **1.66** | - | 1.01 | 0.17 | 0.15 | 0.15 | 0.18 | 6.68 | - | 2.24 |
| Bcor1 | 0.42 | 0.41 | 0.43 | **2.88** | 2.47 | 1.47 | 0.41 | 0.36 | 0.24 | 0.40 | 4.15 | 4.23 | 2.01 |
| Bint1 | 0.38 | 0.44 | 0.51 | **3.64** | 2.76 | 1.59 | 0.61 | 0.58 | 0.32 | 0.53 | 4.87 | 4.65 | 2.06 |
| Bint2 | 0.44 | 0.29 | 0.59 | **3.40** | 2.57 | 1.59 | 0.52 | 0.58 | 0.21 | 0.50 | 4.84 | 4.30 | 1.67 |
| Bnan1 | 0.19 | 0.27 | 0.05 | **1.39** | - | 0.85 | 0.41 | 0.08 | 0.05 | 0.00 | 4.02 | - | 1.99 |
| Bnan2 | 0.11 | 0.25 | 0.30 | **1.27** | - | 0.76 | 0.40 | 0.04 | 0.05 | 0.03 | 4.45 | - | 1.47 |
| Ppro1 | 0.50 | 0.09 | 0.00 | **0.36** | - | 0.00 | 0.29 | 0.02 | 0.05 | 0.00 | 6.08 | - | 1.36 |
| Pweb1 | 0.57 | 0.07 | 0.00 | **0.46** | - | 0.00 | 0.31 | 0.01 | 0.13 | 0.00 | 6.31 | - | 1.46 |
| Ppat1 | 0.82 | 0.16 | 0.02 | **0.80** | 0.66 | 0.04 | 0.47 | 0.16 | 0.13 | 0.00 | 6.14 | 6.04 | 1.43 |

**Table S2:** Genome proportion [%] of different satDNAs (and tandem repeats) across *Beta* and *Patellifolia* species. Abbreviations as in Table 1; accession 1 of the respective species.

| **satDNA proportion [%]** | beetSat01-pBV | beetSat02-pEV | beetSat03-pTS5 | beetSat04-pTS4.1 | beetSat05 | beetSat06-pAv34 | beetSat07-pHC8 | beetSat08 | beetSat09 | beetSat10-pRN | beetSat11 | beetSat12 | beetSat13 | beetSat14 | beetSat15 | beetSat16 | beetSat17 | beetSat18 | beetSat19-Tantalos |
|---|---|---|---|---|---|---|---|---|---|---|---|---|---|---|---|---|---|---|---|
| Bvul | 8.02 | 4.99 | 0.00 | 0.00 | 0.00 | 0.24 | 0.18 | 0.06 | 0.06 | 0.03 | 0.01 | 0.03 | 0.00 | 0.00 | 0.00 | 0.00 | 0.01 | 0.00 | 0.16 |
| Bada | 8.17 | 3.51 | 0.00 | 0.00 | 0.00 | 0.18 | 0.24 | 0.13 | 0.05 | 0.03 | 0.00 | 0.03 | 0.00 | 0.00 | 0.00 | 0.00 | 0.01 | 0.00 | 0.17 |
| Bmar | 3.12 | 3.90 | 0.00 | 0.00 | 0.00 | 0.11 | 0.27 | 0.06 | 0.05 | 0.02 | 0.00 | 0.05 | 0.00 | 0.00 | 0.00 | 0.00 | 0.01 | 0.00 | 0.28 |
| Bptu | 3.90 | 3.92 | 0.00 | 0.00 | 0.00 | 0.18 | 0.27 | 0.07 | 0.07 | 0.02 | 0.00 | 0.03 | 0.00 | 0.00 | 0.00 | 0.00 | 0.01 | 0.01 | 0.22 |
| Bmca | 2.12 | 3.72 | 0.00 | 0.00 | 0.00 | 0.26 | 0.25 | 0.04 | 0.07 | 0.02 | 0.00 | 0.04 | 0.00 | 0.00 | 0.00 | 0.00 | 0.01 | 0.00 | 0.23 |
| Blom | 0.01 | 0.01 | 0.00 | 0.00 | 0.00 | 0.37 | 0.34 | 0.53 | 0.97 | 0.34 | 0.20 | 0.06 | 0.00 | 0.13 | 0.45 | 0.00 | 0.04 | 0.00 | 0.01 |
| Bmrh | 0.01 | 0.54 | 0.00 | 0.00 | 0.00 | 0.78 | 0.47 | 0.53 | 1.45 | 0.76 | 0.38 | 0.40 | 4.06 | 0.12 | 0.43 | 0.16 | 0.06 | 0.00 | 0.03 |
| Bcor | 0.01 | 0.24 | 0.00 | 0.00 | 0.00 | 0.61 | 0.31 | 0.24 | 0.41 | 0.69 | 0.18 | 0.35 | 0.66 | 0.06 | 0.08 | 0.02 | 0.14 | 0.00 | 0.03 |
| Bint | 0.01 | 0.16 | 0.00 | 0.00 | 0.00 | 0.57 | 0.37 | 0.27 | 0.86 | 0.65 | 0.30 | 0.15 | 0.36 | 0.08 | 0.15 | 0.03 | 0.10 | 0.09 | 0.02 |
| Bnan | 0.01 | 0.00 | 0.00 | 0.00 | 0.00 | 0.01 | 0.07 | 0.08 | 0.10 | 1.45 | 0.95 | 0.11 | 0.01 | 0.01 | 0.00 | 0.00 | 0.48 | 0.00 | 0.01 |
| Ppro | 0.00 | 0.84 | 2.69 | 0.50 | 1.75 | 0.30 | 0.00 | 0.00 | 0.00 | 0.00 | 0.00 | 0.00 | 0.00 | 0.00 | 0.00 | 0.00 | 0.00 | 0.00 | 0.00 |
| Pweb | 0.01 | 1.02 | 1.73 | 0.66 | 2.14 | 0.34 | 0.00 | 0.00 | 0.00 | 0.00 | 0.00 | 0.00 | 0.00 | 0.00 | 0.00 | 0.00 | 0.00 | 0.00 | 0.00 |
| Ppat | 0.00 | 0.67 | 1.80 | 0.49 | 2.80 | 0.26 | 0.00 | 0.00 | 0.00 | 0.00 | 0.00 | 0.00 | 0.00 | 0.00 | 0.00 | 0.00 | 0.00 | 0.00 | 0.00 |

**Table S3:** Primer sequences for the amplification of beet-specific satellite DNAs. Whereas genomic DNA of *B. corolliflora* was used for the amplification of beetSat15, beetSat17 was amplified using genomic DNA of *B. nana*.

| primer name | forward sequence 5’-3’ | reverse sequence 3’-5’ | annealing temp. [°C] |
|---|---|---|---|
| beetSat15 | CTC GAG TAA GTT ATT ATG TAA TTG | TTG ATT AAC TCA TTC GTT CAA TTC | 54.0 |
| beetSat17 | CAA GGG GCT CAT TAC TTA GC | GGG TAT TGA GGT ACT TAG CC | 57.3 |

**Data S1:** beetSat nucleotide sequences derived as RepeatExplorer2 consensuses from the comparative analysis (start/end of the monomers are arbitrary).

```
>beetSat01-pBV_consensusmonomer_296nt
ACCTGATTTGAGGGAATTAGACTCACAAACATAATAGGACCCCTATATCATTCAACAACTAA
AAAAAAGAATTATAAAGCAAATGATGTGTTTGTGGATAGTAAAATCAATAACAAGTCTCAAA
ACACCCGAATCTCACTTGTACTTGTTGAAAGATTCGATCTTTATATTGAATAGGCTTTGTTG
TTAGCTCGAGAACTAGAGGATCTTTCATTAACCTCTATTAGGAGATGCACACAAACCTCAAT
TTGTATATTCAATGCAATGCACTCGAATTGTTAAATTAAAACACACAC
>beetSat02-pEV_Beta_consensusmonomer_159nt
TAAGGGGTTCGTGAGCTCAGAAGCTGAATTCGTTGAAGTCCGATTTTTCGCGTCTGTTTCGT
GCTCATCTTGGCCCAAAATCGTAATTTGGGCCTAACTCCTAGCTCGGACATCAGAATTGAGC
GATTCAAAAGCCTGAAATGTTTTTTTTACCATTTC
>beetSat02-pEV_Corollinae_consensusmonomer_158nt
TTTGGGCCCAGAGCAACGTGGAACCGACGCGAAAAATCGGACTTCAATTTTTTCAGCTTCTG
ACCTCACGAACCCTTTAGAAATGGTAAAAAAACGCATTTAGGCGAAAGAATGACTCAAAAAT
GTTATCCGAGCTAGGAGTTAGGCCCAAAACACGT
>beetSat02-pEV_Patellifolia_consensusmonomer_157nt
ATGTCAGGCTTTGGAATCGCTCAAAAATGTTATCCGAGCTAGGAGTTAGGCCAAAAATACGA
TTTGGGCCCCCACGAACAAAGAACCGACGCGAAAAATCGGACTTCAATATTTTCAGCTTCTG
AGCGCTCGAACCCCTTAGAAATGGTAAAAAAAC
>beetSat03-pTS5_consensusmonomer_158nt
TCGCCTAAGAGACTATGACGGTTTTACCCTTTGATTTGAAATGAGTTTGATCCAAGGGCTTC
ATATGCTTTAAATATATCTAATACCTATTCAAGGAGTCAAAAATATTTGGTAATTAAGCACA
ACAAATGATTTGAAAGGTGTTCATACACCACAAA
>beetSat04-pTS4.1_consensusmonomer_312nt
CCCGGCCACGTCCCGGGGCCCGGAGTCGGATCTTGCCCAAAAAATTTTTAGGGCCTCCTTGG
GCCATATGAGGCCCTTTGGAACCTTAACATGCCCCAAAACCGGGCACCGGATGCCTTCCAAT
CCCACCTCCGTAATTCCGACCAATACCGATAACCGACTCGGGAGGCCATTTAGAGGCCCTTT
TGGGGCTAGAAGCCCCCCAAAATTTACAGAGACCCTATTGGGACCCCCGGGAAGGCACCCGT
CAAAAAAAATCACCCTGCCCGAAAATGTTCCGATTTACCCGTTTTCGTGGACTATTACTCAC
GC
>beetSat05_consensusmonomer_164nt
TTTCCATCAACGTAGACTGAGTGCCGTTAAGGAGCATATGTGACTCAAAACCACCTCATACG
ACGGCCGTTTTGCCCCATATTTGGTATTCCATATTTTTTTGGTCATTGAAACCTACCACAAA
AATTTCGTCTCAATCTGGAAACTTTTTTTTGAAACCTTTT
```

>beetSat06-pAv34_consensusmonomer_358nt
TCACCGAACATAGAGATTTTAAAGAATTGTTGAAATCTTTAGAAAAATGGCGTCGGAATGAA
CTTCGACCGTCTAGAAATCAATGCACCGGAACCCTAAGTCTACTCGGGGACCAGAAGAGGCA
TTCTTAGTTTGATTTTATGAAATCTCCGAAATAACACTCAGGCGTGTACCCCGGTTCACTAA
CTCGGAGGATTTTTGAAAATATGTTTGATTTCTCAGAAAATGGCACTGTAAAGCACATTTGA
CCGAAAGGGCCCCAAATAGTGAAAGCCAAAGTCTTCCAAAGGTTTCGTGTTGCCTTTTTAGG
ATGGTTTTAAGTAATCACGAAAAATGCCTCACAGGCGTGTACCGGGGG
>beetSat07-pHC8_consensusmonomer_160nt
TTGTTGGTTGCATTTCGGGAACGAAAAGGTCATTTTAGGCCTATTTTTGCCTTTTCGCTTCC
AAATCGTACTTATCGAGTCATTTAACCCCAAATTCTTTTTGAACATGATTATATGAGGTATA
ATGTAACTATTGTGTAGATTTTTAGGCTTGCATTTG
>beetSat08_consensusmonomer_171nt
ATATTAGCCTATATTTGGTTCTAACTTTCCTATTTTGTGTATTTTCAATCAAAACAACTTAT
AATTACATTAAACTAACTATATATATGATGTTTGAGGTGTTTATAACTTAAACTAGCATCAA
ACTACTCGTATTCATTCATTTTCAAGTAAAATGGCCTAAAATGGCCT
>beetSat09_consensusmonomer_160nt
TATGAGTGGACTTTAATTGACTCTTATAGTTATATATGTACCTATTGGAACTATATATGACC
TAACATGTGGCTAAATGCGGGAAACTAAGTGAATTGAGCTTCAAAGGCCAAAATCGGCCCGA
AATGGCCTTCATTTACCCAAAATGGGTTTCAAAGCA
>beetSat10-pRN_consensusmonomer_214nt
ACAATCCATTCCCATTACTCATCAAAGCAATCCTATGGGGATTGGACAATCATGCCACTGGG
TCCATTAGGCCCAATACCTCACCAAACCCCATTGACTTGAGTTTTGGGCCCCCATTGTCCCT
CCTTGTACTAAAAAGCCCAAATTAACCCCATATCATATGCTTTTAGGTAATTACCAAACATG
TTTTGGGCATTACCAAACCTATCCAATC
>beetSat11_consensusmonomer_157nt
CGGACTTGAGGAATCTGTTGAGCATGGAAAAATGCCACTTTAGGCGCAAGAATCAATTAGAA
ATGTTTTTCGAGATAAGAGTTATGGGAAAAATACTTTTCACGTTCAATGTAGGGCACTACTT
GTACGAAAAATCCAACTTCATTTTTTTCCCATT
>beetSat12_consensusmonomer_518nt
TTTTGATCGAGTAGATTTCGAGTAACTTGCAAGTGTCTCGTGTTAAAAGCAACCCTAAGGTA
GAAAATATGCCAAATTTAGGTATTTTGTATAGATAATTGGGTATTCGAGCTTCTCACGAGTT
TTCGAGCCGAGTATCGAGTTGCTTAACTCGACTCGACAACATCTCGAGTCGAACTCAAGTTC
GATTCGAGCTCACTCGAGTCGAGCTTTGACCGAGTCGATCTCGAGTAGCCCGCAAGCCGTCT
CGTCTCATTTACACCTCTACCCTTGCATTGTTACTTTGGTATCATTGAAATGATACTTTGGT
ACCATTAAAAGGGTAATTTGGTACACTTGCATTATTTCTTTGGTATTATTCAAATGATACTT
TGGTACCATTAAAACGGTAATTTGGTACCCTTGCATAGGGTGTAAATGATGCGAGATGAGTT
CGAGTTTGAGCTAGCTCGACTCGACTTGTTAGCATTTCGAGTCGATCTTTGACCGAGTATTG
ACCGAGTTTTGACCGAGTCGAG
>beetSat13_consensusdimer_79nt
TTTGATTCAATTAGCTTTGTTTGACTTCATTTGACTTTCATTTGATTCAATTAGCTTTGTTG
AATGCATTTGAGTTTCA
>beetSat14_consensusmonomer_162nt
TTTGGGAAGTGAAATAGCTCAAATTCACAAAAATCGACCCCGAAACGAACCTAAAAATTGAA
CTTCTAATCTTTTGCACCTAAATATGACATTAACACTTCAACTATGCATAAAGTAAGTGGTG
TGTGTTAGTTTTGCTCCTATATGTAACATGTAGCTAGT
>beetSat15_consensusmonomer_63nt
AATAACTAAATAAAACACTCGAGTAAGTTATTATGTAATTGAATTGAACGAATGAGTTAATC
A
>beetSat16_consensusmonomer_347nt
GTCAGACTCGGATGATGACCTAAATTTCTTTTGCAACGAGAATTTAAAGATTTTGGAATTAG
TATAAAAAAGCATGGGGTTCTCTTCTAGTAGACCCTACTGATTTGCAATTGCATTCACATGA
ATTAATTATGAAACAGTGTTTGCTACTTGCAGTTTACAATACTTGAATGTTTAATCCATGTA

TAAAATTGTACTGTTTTCCTTCTTGTTAAATTTTAAAGGGAAAGCTTGGAAGTTGCTACTTT
GGACTTTGAAGGATTGAATTACATTCGGCACAAATCCAAAATTGTTTTGTTATATTTTAGGT
CTTGACCTGAGTAAATTTTGCGAATTACTTTGTAGCT
>beetSat17_consensusmonomer_112nt
ATGGAAGGGCTCATTAGCCCCCGAGTTGGTGCACACCCTTCTCGGCCTTGCCAGGGGGCTCA
ACACTTAGCCAAAAATTGGCTAAGTACCTTCATGCCCAACTCGGGATCCC
>beetSat18_consensusmonomer_65nt
TCGACGAGGAACGTCCAGACATGGACGTCCTCCTTCTACGAAGGACCCTCGTCCAGAAATCC
TCC
>beetSat19-Tantalos_Zakrzewski_325nt
TGTGACTTGTAACATTGCGCGGGTGCTTGGCACCATTTGCGTTACCTCAAAAAGCCTTTGAA
CACCCCAATTATTCATTTCTCGCGAAATCCAAAATTGCCTCGAAATGAACGTAAAGGCATCC
ACATATTTGTTCCAAGCCACATGACTCCTTTACATTGACCTCCTATGTCCCTAGGAGGCATC
CCGTGCCATTTGGAGCTCGGGCAACGGGAAAGTCCGAAAGCGTGTATAATCTTCAATTTTAG
TTGTTTTTGGGGAATTTTTGGACTACTTCTTCAGGCCCGGTCATATTTTTCTTTCGAAACAT
TCCTAGGAGTGCCGA

**Data S2:** beetSat sequences used as FISH probes (the beetSat05 probes were synthesized as EXTREmer oligonucleotides).

>beetSat01-pBV_probe_Z22849.1
GGATCCACACAAACCTCTATTTGTAAGTTCAATGCAACGAACTCGAATTGTTAAATTAAAAT
ACACAAACAGTTGATTTGAGGGAATTAGACTTACAAACATAATAGGACCCCCATATCATTAA
AAAACAAAAAAAAAAAATTATAAAGCAACTGATGTGTTTGTGGATAGTAAAATCAATAACAA
GTCTCAGAACACTCGAATCTCACTTCTACTTGTTGAAAGATTCGATCTTTATATGAAATAGG
GCATAAAGATTGAATCTTTATATTGAATAGGCTTTGTTGTTTGCTCGAGAACTAGAGGATCT
TTCATTAACCTCTATTA
>beetSat02-pEV_probe_OY726583.1
GAATTCGTTAAAGTCCGATTTTTTGCGTCTGTTTCGTGCTCATCTTGGCCCAAACTCGTAAT
TTAGGCCTAAATCCTAGCTCGGACGTCAGAACTGATCGAACCAAAAGCATGAAATTTTTTTT
TACCAT
TTCTAAGGGGTTCGTGAGCTCATAAATTGAATTCGTTAAAGTCCGATTTTTTGCGTCTGTTT
CGTGCTCATCTTGGCCCAAACTCGTAATTTAGGCCTAAATCCTAGCTCGGACGTCAGAACTG
ATCGAA CCAAAAGCATGAAATTTTTTTTTACCATTTCTAAGGGGTTCGTGAGCTCATAAATT
>beetSat03-pTS5_probe_Z50809.1
GATCCAAGGCTTAGTATGCTTTACATATACCTAATACCTATTAAAGGAGTCAAAAACAATAG
GTAATTAAGCACATCAAATGATTTGAAAGGTGTTTCATACACAACAAATCTCGTAAGATACT
ATGATAGTTTTAACCTTTGATTTGAAATGAGTTT
>beetSat04-pTS4.1_probe_Z50808.1
GATCATGTCCAAAAATATTTTAGGGCCTCCTTGGGCCAAATGACGCCCTTTGGAACCTTAAC
ATGCCGAAAATCGGGTACCGGATACCTTCCAATCCCACCTCCGTAATTCCGACCAATATCGA
TAACCGACTCGGGAGGCCATTTAGAGACTCTTTTGGGGCTTGAAGCCCCCCAAAATTTACAG
AGACCCTATGGGGACCCCCGGGAAGGCACATGTGAAAAAAAATTACCCTGCCCAAAAATGTT
CCGATTTACCCGTTTTTGGTGGACTATTACTAACGCCCCGGCCACGACCCAGGGTCCGGAGT
TG
>beetSat05_Oligo-fw
TTTCCATCAACGTAGACTGAGTGCCGTTAAGGAGCATATGTGACTCAAAACCACCTCATACG
ACGGCCGTTTTGCCCCATATTTGGTATTCCATATTTTTTTGGTCATTGAAACCTACCACAAA
AATTTCGTCTCAATCTGGAAACTTTTTTTTGAAACCTTTTA

```
>beetSat05_Oligo-rv
AAAAGGTTTCAAAAAAAAGTTTCCAGATTGAGACGAAATTTTTGTGGTAGGTTTCAATGACC
AAAAAAATATGGAATACCAAATATGGGGCAAAACGGCCGTCGTATGAGGTGGTTTTGAGTCA
CATATGCTCCTTAACGGCACTCAGTCTACGTTGATGGAAAA
>beetSat07-pHC8_probe_AJ243337.1
GGCCAATTTTTGCCTTTTCGCGTCCCTTTCGTACTTATCGAGTCATTTAACCCCAAATTATT
TTTGAACATGATTATATGAGGTATAATGTAACTATTGTGTAGATTTTTAGGCTTGTATATGT
TTTTGGTTGAATTTCGGGAACGAAAAGGTCATTTTAG
>beetSat08_probe_OY726584.1
CTAGTTTAAGTTGTAAACACCTCAAACATCATATATATAGTTAGTTTAATGTGATTGTAAGT
TGTTTTGATTGAAAATACACAAAATAGGAAAGTTAGAACCAAATATAGGCTAATATAGGCCA
TTTTAGGCCATTTCACTTGAAAATGAATGAATACGATTAGGTTGGTGCTAGTTTAAGTTATA
AACACCTCAAACATTATA
>beetSat10-pRN_probe_Z69354.1
ACTAAAAAGCCCAAATTAACTCCATATCATATGCTTTTTAGTAATTACCAAACATGTTTTGG
CCATGACCAAACCTACTAAGTGGAGGATGTTCTCCCAATCACAATCCATTCCCATTACTCAT
CAAAGCAATCCTATGGGGATTGGACAATCATGCCACTGGGTCCATTAGTCCCAATACCTCTC
CAAACCCCATTGACTTCAGTTTTGGGCCCCATTGTCCATCCTTGT
>beetSat13_probe_LR215734.1
GTTTGACTTTCATTTGATTCAATTAGCTTTGTTGGATGAATTTGACTTTCATTTAATTCAAA
AAGCTTTGTTTGAATGTGTTTGTCATTCATTTGATTCAATTGGCTTTGTTCGAATGTTTTTG
ACTTTCATTTGATTCAATTAGCTTTGTAGAATGCATTTGACTTTCATTTAATTCAAATAGCT
TTGTTTGTGAAGTTTGTTTCGA
>beetSat15_probe_OY726585.1
ATTTGATTAACTCATTCGTTCAATTCAATTACATAATAACTTACTCGAGTATTTTATTTAGT
TATTTGATTAACTCATTCGTTCAATTCAATTATATTATAACTTACTCAAGTGTTTTATTTAG
CTATTCGATTAACTCATTTCGTTCAATTCAATTACATAATAACTTACTCGAGTGTTTTATTT
AGTTATTTGATTAACTCATTCGTTCAATTCAATTGCATAATAACTTACTCGAGTGTTTTATT
TAGTTATTTGATTAACTCATTTCTTCAATTCAATTACATAATAACTTACTCGAGA
>beetSat17_probe_OY726586.1
CAAGGGGCTCATTACTTAGCTAAAAAATGGCTGAGTACTTCAATACCTAACTCGGGATCCTC
ATAGAAGGGGCTCATTGGCCCCCGAGTTGGTGCATCACCCTTCTTGGCCTTGCCAAGGGGCT
CATTACTTAGCCAAAAAATGGCTAAGTACTTCAATACCCAACTCGGGATCCTCATAGGAAGG
GCTCATTAGCCCCCGAGTTGGTGAATCACCCCTCTTGGCCTTGCCAACGGGCTCATTACTTA
GCTAAAAAATGGCTAAGTACCTCAATACCCA
```
